# Dissociation of attentional state and behavioral outcome using local field potentials

**DOI:** 10.1101/2023.08.05.552102

**Authors:** Surya S Prakash, J Patrick Mayo, Supratim Ray

## Abstract

Successful behavior depends on attentional state and other factors related to decision-making, which may modulate neuronal activity differently. Here, we investigated whether attentional state and behavioral outcome (i.e., whether a target is detected or missed) are distinguishable using the power and phase of local field potential (LFP) recorded bilaterally from area V4 of monkeys performing a cued visual attention task. To link each trial’s outcome to pairwise measures of attention that are typically averaged across trials, we used several methods to obtain single-trial estimates of spike count correlation and phase consistency. Surprisingly, while attentional location was best discriminated using gamma and high-gamma power, behavioral outcome was best discriminated by alpha power and steady-state visually evoked potential. Power outperformed absolute phase in attentional/behavioral discriminability, although single-trial gamma phase consistency provided reasonably high attentional discriminability. Our results suggest a dissociation between the neuronal mechanisms that regulate attentional focus and behavioral outcome.

## Introduction

Attention facilitates behavior by selectively processing relevant sensory information (Carrasco 2011). Attention improves behavioral performance by modulating neuronal signals such as spike rate (Motter 1993; McAdams and Maunsell 1999) and local field potential (LFP) oscillations in various frequency bands (Fries et al. 2001; Khayat et al. 2010; Prakash et al. 2021). The power in different frequency bands have often been used to decode the focus of attention (Esghaei and Daliri 2014; Tremblay et al. 2015; De Sousa et al. 2021; Prakash et al. 2021). For example, we recently found that high-frequency LFP oscillations (>30 Hz) decode attention better than or on par with spikes when multiple electrodes are used (Prakash et al. 2021). This result contrasts with sensory (Kanth and Ray 2020) and motor decoding(Hwang and Andersen 2013) where multichannel spikes outperform multichannel LFP. To account for these differences in the mechanisms of cognitive vs sensorimotor processing, we proposed that the elevated performance of high-frequency LFPs is due to comparable spatial spreads of LFP and biophysical circuits that underlie attention (Prakash et al. 2022).

Our previous study (Prakash et al. 2021) probed a single aspect of behavior, the attentional state of the animal. Because there is no objective measure of where the monkey’s attention was focused at a particular moment, the decoding may have been more related to which side the attentional cue was given and where the focus of attention was located on average. Specifically, we associated attention (as conventionally done) with cue-in (cue inside the receptive field (RF) of neurons being recorded) and cue-out conditions, which we termed “attention-in” and “attention-out” to be consistent with previous literature (Fries et al. 2001; Khayat et al. 2010). However, another relevant decoding variable that has been extensively studied is the behavioral outcome of a trial (“hit” for a detected target change versus “miss” for an undetected target change). Many studies have reported that the phase of low-frequency oscillations is informative for decoding behavioral outcome (Busch and VanRullen 2010; Harris et al. 2018). However, whether decoding behavioral outcomes based on LFP features is as informative as decoding based on spikes is not known. This evaluation is critical for designing functional brain-computer interfaces and for potentially revealing the neural basis of attention and decision making.

LFP phase consistency has been hypothesized to play a particularly important role in communication between neuronal ensembles and has been shown to be modulated by attention (Bosman et al. 2012; Grothe et al. 2012; Fries 2015). For example, attention enhances the spike-LFP and LFP-LFP phase coherence in the gamma band in macaque visual area V4 (Fries et al. 2001; Buffalo et al. 2011; Prakash et al. 2021). Likewise, behavioral performance measured using reaction time also varies with the absolute LFP phase (Ni et al. 2016), spike-LFP phase coherence (Womelsdorf et al. 2006) and inter-areal LFP-LFP phase consistency (Rohenkohl et al. 2018) in the gamma band. Collectively, a substantial body of work suggests that phase relationship between neuronal population is systematically modulated by both attention and behavioral performance.

Because phase consistency measures are typically calculated across trials (but see Spyropoulos et al. 2024), they may fail to capture the well-established trial-by-trial changes in attentional fluctuations and behavioral performance that we seek to understand (Cohen and Maunsell 2010, 2011; Ghosh and Maunsell 2021; Roy et al. 2021). Thus, existing methods for calculating phase consistency may provide only a coarse estimate of the underlying dynamics of attention or behavioral state. This shortcoming is not limited to LFP measures. Conventional measures of spike count correlation between two neurons—which have been shown to reduce with attention (Cohen and Maunsell 2009; Mitchell et al. 2009; Mayo and Maunsell 2016)—suffer from the same limitation. To address these issues, we used several methods to compute pairwise metrics such as spike count correlation, LFP power correlation, and field-field coherence for single trials. We tested the ability of spikes, LFP power, LFP phase, and pairwise metrics in visual area V4 to independently decode attention and behavioral outcome of two rhesus macaques performing a standard change detection task.

## Materials and Methods

### Electrophysiological recordings

Data and experimental procedures used in this study are described in our previous studies (Mayo and Maunsell 2016; Prakash et al. 2021). We outline the details briefly here. All the animal procedures were approved by the Institutional Animal Care and Use Committee of Harvard Medical School. Two adult male rhesus monkeys (*Macaca mulatta*) were surgically implanted with a titanium headpost and scleral eye coil before training on the task. After training, each monkey was implanted with two 6 × 8 (48) microelectrode Utah arrays (Blackrock Microsystems), one in visual area V4 of each hemisphere. Electrodes were 1 mm long and 400 µm apart. The impedances of the electrodes were in the range of 0.2 – 1MΩ at 1 KHz.

The raw signals from 96 channels were recorded using a 128-channel Cerebus Neural Signal Processor (Blackrock Microsystems). Single and multi-unit spiking activity were obtained by filtering the raw signal between 250 Hz (4^th^ order digital Butterworth filter) and 7.5 kHz (3^rd^ order analog Butterworth filter) and subsequently subjected to a voltage amplitude threshold and sorted offline using spike sorting software (Plexon). LFP signal was obtained by filtering the raw signal between 0.3 Hz (1^st^ order analog Butterworth filter) and 2.5 kHz (4^th^ order digital Butterworth filter), sampled at 2 kHz and digitized at 16-bit resolution. The possible aliasing caused by inadvertently setting the low pass LFP filter cutoff at a frequency that was higher than the sampling frequency had negligible effect on the LFP (Prakash et al. 2021).

### Behavioral task

Monkeys were trained to perform an orientation change detection task (Fig. 1). A trial began with the appearance of a small fixation spot at the center of the monitor, and the monkeys held their gaze within a 1.8° square window centered on the fixation spot throughout the trial. Two full contrast Gabor stimuli were presented simultaneously and synchronously on a CRT monitor (100 Hz frame rate, 1024 × 768 pixels) over a uniform gray background, positioned 57 cm from the monkeys. Each Gabor was sinusoidally counterphased at 10 Hz and its size, location, spatial frequency, and initial orientation were optimized for a randomly selected recording channel in each hemisphere during each session. At a random time chosen from an exponential distribution (mean: 3000 ms; range: 500 – 5500 ms) when the contrast of both the stimuli were at 0%, one of the stimuli changed its orientation. Monkeys were rewarded for making a saccade to the changed stimulus between 100 and 550 ms after the change. Eye position was sampled at 200 Hz using the scleral eye coil technique (Judge et al. 1980).

**Figure 1:**
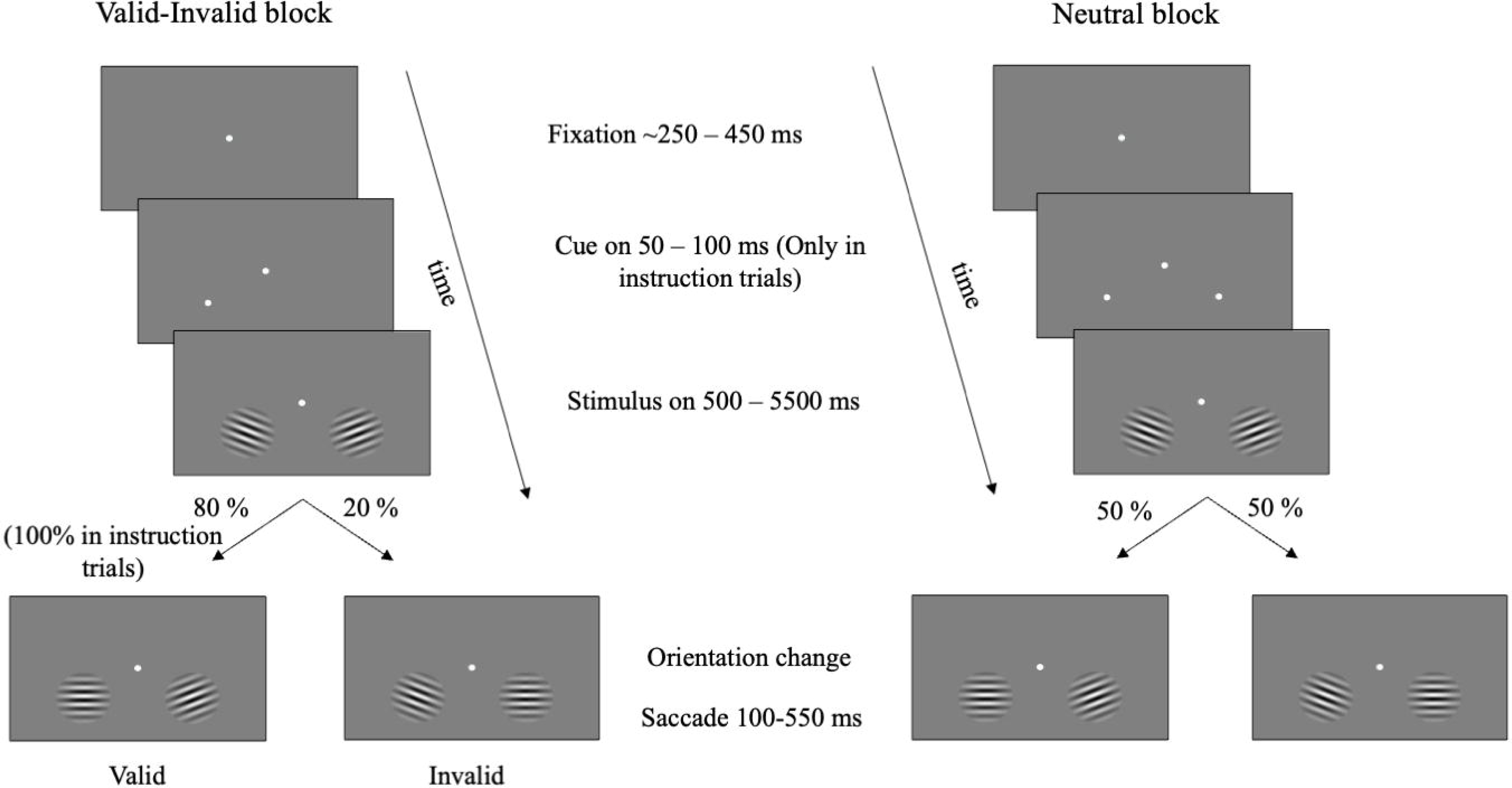
Schematic of orientation change detection task. Valid-invalid block (left panel): Task began with the appearance of a central fixation spot. Attention was cued in blocks using an initial set of “instruction trials” where the monkeys were explicitly cued to attend to a location (cued left in the depiction) by briefly flashing a white dot for 50 – 100 ms which indicated the target location with 100% certainty in the instruction trial and most likely location for rest of the block. Two gabor stimuli counterphasing at 10 Hz appeared simultaneously and synchronously on the screen. At an unsignalled time between 500 to 5500 ms after the stimulus onset one of the gabor changed its orientation (target) and monkey was rewarded if it made a saccade to the location of orientation change between 100 – 550 ms after the target onset. Target appeared at the cued location with 80% probability (valid cue) and at uncued location with 20% probability (invalid cue). Monkeys were rewarded for detecting the target at either of the locations. Once the monkeys performed four instruction trials correctly then the explicit cue was no longer presented, and the target appeared probabilistically. Neutral block (right panel): Unlike the valid-invalid block, in neutral block, the monkeys were cued by briefly flashing the white spot at both the locations simultaneously during the instruction trials and the target could occur at either of the two locations with 50 % probability. The rest of the task structure is same as that of valid-invalid block

Monkeys were cued to attend to one (valid-invalid condition) or both stimuli (neutral condition) in blocks of ∼50 trials, each beginning with four correct instruction trials in which a small white spot appeared briefly for 50 to 100 ms indicating the location(s) of the change (target). In valid-invalid blocks, the target appeared at the cued location with 80% probability (valid cue) and 20% probability at the uncued location (invalid cue). In neutral blocks, the target appeared at either location with 50% probability. Five percent of the total trials were catch trials in which no change occurred, and monkeys were rewarded for maintaining the fixation throughout the trial. Catch and instruction trials were not included in the analysis. Six different orientation change magnitudes were used for valid and neutral cue conditions and only the second and third smallest orientation change were used to probe attention in the invalid condition in each recording session.

## Data analysis

We analyzed the data from 25 sessions of two monkeys (Monkey A: 13 sessions; Monkey W: 12 sessions). We restricted our analysis to trials with either the second or third smallest orientation change in which the behavioral performance was close to 50% to obtain a comparable number of trials between hit and miss conditions. Note that the magnitude of orientation change can be different for different cue types and locations. In addition, we considered data for a particular cue condition of a given session only if there were more than 10 trials in each of their four conditions. Consequently, 22 out of 25 sessions met the criteria for valid and neutral cue but none for invalid cue. Hence, we only analyzed the data of valid and neutral cue conditions.

### Mean matching of target onset time distribution

We matched the target onset time distribution of hit and miss trials of each attention condition to remove the artifactual drift introduced due to the difference in the number of early target onset (<750 ms) trials. We sorted the hit and miss trials separately based on the target onset times (500-5500 ms) and divided them into bins of 250 ms and randomly chose an equal number of trials from both the conditions in each bin. We performed the analysis (Fig. 2 onwards) on the mean matched data and computed all the neural measures. The process was repeated 50 times and then averaged across iterations.

**Figure 2:**
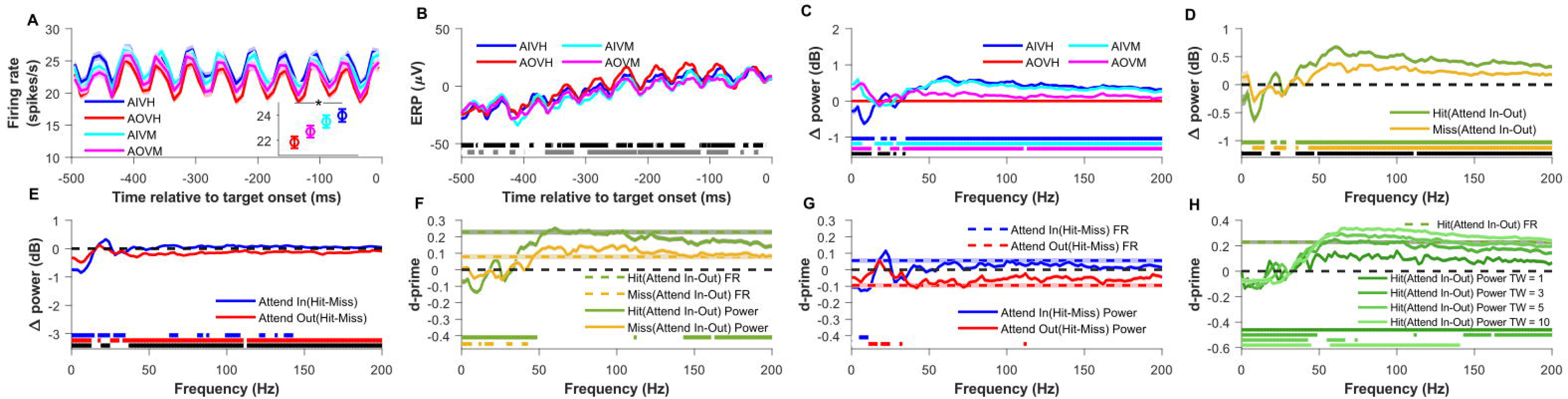
Comparison of firing rate (FR) and local field potential (LFP) power across validly cued attention and behavioral conditions for matched target onset time distributions. (A) Mean peri-stimulus time histogram (PSTH) relative to the target onset time for the conditions in which attention was validly cued into or outside the receptive field of a neuron and the subject either detected (hit) or missed the target, namely, Attend-In Valid Hit (AIVH; blue), Attend-Out Valid Hit (AOVH; red), Attend-in Valid Miss (AIVM; cyan) and Attend-Out Valid Miss (AOVM; magenta). Inset shows the mean firing rate over the same time period as PSTH for the four conditions. The mean is first taken across 643 electrodes recorded across 21 sessions in two monkeys and then averaged across 50 bootstrap iterations of target onset time matched trials selection. Shaded lines and error bars indicate the bootstrap mean of s.e.m across the 643 electrodes. Only sessions in which at least 10 stimulus repeats were available for every condition were used for analysis. Asterisk in the inset indicate that mean firing rate of AIVH is significantly higher than AOVH (Wilcoxon rank sum test; Bonferroni corrected p<0.05). (B) Mean event-related potential (ERP) for the four conditions. Horizontal black patches at the bottom indicate the time values during which the ERP of AIVH and AIVM were significantly different (Wilcoxon rank sum test; Benjamini-Hochberg FDR controlled p<0.05) and grey patches indicate the time values during which ERP of AOVH and AOVM were significantly different (Wilcoxon rank sum test; Benjamini-Hochberg FDR controlled p<0.05). (C) Mean change in power spectral density (PSD) in decibels for all the validly cued conditions relative to Attend-out Valid Hit (AOVH) condition. The horizontal blue, cyan and magenta patches at the bottom indicate the frequencies at which the change in PSD relative to AOVH condition is significantly different from zero of the conditions represented by their corresponding color (Wilcoxon signed-rank test, Benjamini-Yekutieli FDR controlled p<0.05 under unknown dependency). Black horizontal patches indicate the frequencies at which the change in PSD of AIVH (blue trace) and AIVM (cyan trace) are significantly different from each other (Wilcoxon rank-sum test, Benjamini-Yekutieli FDR controlled p<0.05 under unknown dependency). (D) Mean change in LFP power spectral density (PSD) in decibels between attend-in and attend-out conditions for hit (AIVH and AOVH; green) and miss (AIVM and AOVM; yellow) conditions. The horizontal dashed black line marks the zero of the y-axis. Horizontal green and yellow patch at the bottom indicate the frequencies at which the change in PSD between attention conditions are significantly greater than zero for hits and miss conditions respectively (Wilcoxon signed-rank test, Benjamini-Yekutieli FDR controlled p<0.05 under unknown dependency). Horizontal black patch at the bottom indicate the frequencies at which the change in PSD between attention conditions for hit and miss condition are significantly different from each other (Wilcoxon rank-sum test, Benjamini-Yekutieli FDR controlled p<0.05 under unknown dependency). (E) Mean change in power spectral density between hit and miss condition for Attend-In (AIVH and AIVM; blue) and Attend-Out (AOVH and AOVM; red) conditions. The horizontal dashed black line marks the zero of the y-axis. Horizontal blue and red patch at the bottom indicate the frequencies at which the change in PSD between behavioral conditions are significantly greater than zero for attend-in and attend-out conditions respectively (Wilcoxon signed-rank test, Benjamini-Yekutieli FDR controlled p<0.05 under unknown dependency). Horizontal black patch at the bottom indicates the frequencies at which the change in PSD between behavioral conditions for attend-in and attend-out condition are significantly different from each other (Wilcoxon rank-sum test, Benjamini-Yekutieli FDR controlled p<0.05 under unknown dependency). (F) Mean d-prime (the ratio of the mean difference and pooled standard deviation of two conditions) of LFP power (solid lines) and firing rate (dashed color lines) between attend-in and attend-out conditions for hit (green) and miss (yellow) conditions. The horizontal dashed black line marks the zero of the y-axis. Horizontal green and yellow patches at the bottom indicate the frequencies at which the d-prime of LFP power is significantly different from the d-prime of firing rate for hit and miss conditions respectively (Wilcoxon rank-sum test, Benjamini-Yekutieli FDR controlled p<0.05 under unknown dependency). Negative d-prime values were multiplied by -1 before performing the significance test because we were interested in the differences in magnitude of d-prime. (G) Mean d-prime of LFP power (solid lines) and firing rate (dashed color lines) between hit and miss condition for Attend-In (blue) and Attend-out (red) conditions. The horizontal dashed black line marks the zero of the y-axis. Horizontal blue and red patches at the bottom indicate the frequencies at which the d-prime of LFP power is significantly different from the d-prime of firing rate for attend-in and attend-out conditions respectively (Wilcoxon rank-sum test, Benjamini-Yekutieli FDR controlled p<0.05 under unknown dependency). Negative d-prime values were multiplied by -1 before performing the significance test as in (F). (H) Mean d-prime of LFP power between attend-in and attend-out conditions for hit condition where LFP power is estimated by Multitaper method using time-frequency bandwidth product (TW) of 1 (darkest green), 3 (dark green), 5 (light green) and 10 (lightest green). The dashed green line indicates the d-prime of firing rate between the same conditions. The horizontal dashed black line marks the zero of the y-axis. Horizontal colored patches at the bottom indicate the frequencies at which the d-prime of LFP power estimated using different TW is significantly different from the d-prime of firing rate (Wilcoxon rank-sum test, Benjamini-Yekutieli FDR controlled p<0.05 under unknown dependency). Negative d-prime values were multiplied by -1 before performing the significance test as in (G and F).

### Spiking and LFP analysis

We analyzed the spikes and LFP data between 500 to 0 ms before the target (orientation change) onset to obtain the following measures.

### Single electrode measures

#### Trial-wise measures

##### Peristimulus time histogram (PSTH)

PSTH was obtained by counting spikes in non-overlapping 10 ms bins for all the trials and converting them into firing rate which were then averaged across trials.

##### Evoked response potential (ERP)

ERP is obtained by trial averaging the LFP signal in the analysis period.

##### Single-electrode pairwise phase consistency (PPC)

Traditionally, PPC quantifies the consistency of phase difference between two signals, like spike and LFP from the same electrode or spike-LFP or LFP-LFP from two different electrodes (Vinck et al. 2010). In single-electrode PPC, we simply used the absolute phase rather than phase difference. We estimated five phase values using the Multitaper method with five Slepian tapers (Jarvis and Mitra 2001) using Chronux toolbox (RRID:SCR_005547) (Bokil et al. 2010) in MATLAB (RRID:SCR_001622). Then, we calculated the PPC for the phase estimate of each taper and averaged the PPC values across tapers to obtain the PPC value for each electrode.

### Single trial measures

#### Power spectral density (PSD)

We estimated the power of the LFP signal in 0 to 200 Hz frequency range for each trial with 2 Hz resolution using the Multitaper method with five Slepian tapers.

#### Phase

Phase was estimated by taking the Fourier transform of the LFP signal. To account for the circular nature of the phase data, we used the sine component of the phase angle as the measure.

### Electrode pairwise measure

#### Trial-wise measure

##### Firing rate/ LFP power correlation

Pearson correlation between the firing rates or LFP powers of pairs of electrodes were computed across trials.

##### LFP-LFP PPC

The phase relationship between the LFP signals of pairs of electrodes was quantified using PPC, an unbiased estimate of square of the phase lock value measuring the consistency of the phase difference of LFP signals between two electrodes across pairs of trials (Vinck et al. 2010). We obtained five phase angles at each frequency for each electrode using Multitaper method with five Slepian tapers and then computed the PPC of taper averaged phase difference between two electrodes across pairs of trials.

#### Single trial measure

Traditionally, electrode pairwise measures like correlation and PPC are measured across trials. But we needed a single trial measure of these pairwise measures to predict the attention and behavioral state of the monkeys on each trial. Hence, we developed the following two methods to estimate the correlation and phase consistency of a single trial.

1. Binning: We divided spiking and LFP data of 500 ms length into ten non-overlapping segments of 50 ms each. We computed firing rate and LFP power in each segment resulting in ten estimates of firing rate and LFP power with a frequency resolution of 20 Hz. Correlation and phase consistency were computed across these ten estimates. Since this binwise correlation captures the fluctuations over time, factors like adaptation where the activity of the neurons reduce over time can also give rise to spurious correlation. To remove the effects of such factors, we shuffle corrected the correlation values (Smith and Kohn 2008). We calculated binwise correlation values for a given trial with every other non-simultaneous trial and averaged it across all the trial pairs. We then subtracted trial pair averaged correlation from the original binwise correlation of the corresponding trial.
2. Multitaper: Multiple independent estimates of LFP power and phase for a trial can also be obtained using Multitaper method, where a single trial LFP data is multiplied by several orthogonal tapers and then Fourier transformed to obtain several estimates of power (by squaring the amplitude) and phase (see Fig. 4). We used nine tapers to obtain nine independent estimates of power and phase for a trial and computed correlation and phase consistency across them.

In addition to the above two methods, we also calculated single-trial estimates of LFP power correlation and pairwise phase consistency using the Hilbert transform method (Spyropoulos et al. 2024). We band-pass filtered the LFP signal within the analysis period using 4^th^ order Butterworth filter of center frequency starting from 6 Hz with a frequency bandwidth of 10 Hz (i.e., center frequency ± 5Hz) moving in step of 2 Hz up to 200 Hz. The LFP signal filtered around each center frequency was Hilbert transformed to obtain instantaneous amplitude and phase values at each of the 1000 timepoints within the 500 ms analysis window. Instantaneous power was calculated by squaring the amplitude. Finally, correlation in power and phase consistency were calculated across timepoints for every trial and correlation values were shuffle corrected.

### Discriminability using d-prime

We were interested in quantifying the separation in the distributions of the responses for two classes (Attend-In versus Attend-out or Hit versus Miss). Such discriminability is typically quantified using d-prime, which quantifies the difference in the means of the two distributions normalized by their average standard deviation. Here, we used pooled standard deviation to account for unequal number of trials between conditions. The advantage of this measure is that no explicit threshold needs to be assumed for classification. Another equivalent metric is the area under the receiver operating characteristic curve (AUC), which is related to d-prime for normally distributed classes by AUC = 0.5 + erf (d-prime/2)/2, where erf is the error function. Therefore, d-prime values of 0.1, 0.2 and 0.3 (which is about the maximum we obtained in our data) correspond to AUC values of 0.52, 0.55 and 0.58. We used d-primes for discriminability due to their ease of computation and interpretation.

The d-prime metric is given by the formula:

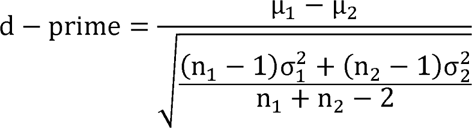

where, µ_1_ = mean across trials of attend/target-in or hit condition

µ_2_ = mean across trials of attend/target-out or miss condition

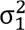 = variance across trials of attend/target-in or hit condition

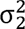 = variance across trials of attend/target-out or miss condition

*n*_1_ = number of trials in attend/target-in or hit condition

*n*_2_ = number of trials in attend/target-out or miss condition

### Discriminability of traditional frequency bands

We compared the discriminability of the neural measures in the following standard frequency bands: alpha (8-12 Hz), gamma (40-80 Hz), high-gamma (120-200 Hz) and SSVEP (18-22 Hz) frequency. We chose SSVEP frequency over a band of frequency rather than a single-frequency point because of frequency smoothing caused by using five tapers. To obtain the power and PPC in each band we averaged them over the frequencies of their respective bands. We excluded the harmonics of SSVEP frequency (multiples of 20) in the gamma and high-gamma band. However, for the binning method where the frequency resolution was 20 Hz, we excluded only 60 Hz. We computed the correlations of the frequency-averaged power.

We could not analyze the behavioral discriminability (hits vs miss) at the population level by combining the activity of all the electrodes like we did previously (Prakash et al. 2021) for attention discriminability using linear discriminant analysis (Fisher 1936) because there were an insufficient number of miss trials. The invertibility of the covariance matrix (required to obtain the weight vector) is guaranteed when the sample size is greater than the dimension (Li et al. 2006). But in our dataset, the number of missed trials (∼20 trials (samples) per session on average after matching the target onset times) was lesser than the number of electrodes (∼30 electrodes (dimension) per session on an average). Although, this limitation can be overcome by regularization (Guo et al. 2007), it underestimates the discriminability (see Supplementary Fig. 2 of Prakash et al. 2021).

### Statistical analysis

We performed Wilcoxon signed-rank test to compare whether the difference between attention and behavioral conditions for each neural measure was different from zero. We used Wilcoxon-rank sum test to compare the attention and behavioral differences across neural measures and discriminability (d-prime) of different neural measures. Negative d-prime values were converted to positive values by multiplying with -1 before performing the significance test since we were comparing only the magnitude. All the p-values are reported after correcting for the multiple comparisons using Bonferroni correction or Benjamini-Hochberg procedure controlling for false discovery rate (FDR) for independent statistical tests and Benjamini-Yekutieli procedure for dependent statistical tests under unknown dependency. Multiple comparison test was done using Multiple testing toolbox (Martínez-Cagigal 2024) in MATLAB. We performed the statistical tests on the bootstrap averaged measures across the electrodes.

## Results

We recorded spike activity and local field potentials (LFPs) using microelectrode arrays implanted in both hemispheres of area V4 in two monkeys. Monkeys performed an attention task in which two Gabor stimuli that counterphased at 10 Hz were presented on a screen and the monkeys were cued to either attend to one or both stimulus locations (see Methods for details) in separate blocks of trials (Fig. 1). In trials where one location was cued, the target (change in the orientation of the Gabor) occurred on the cued side in 80% of trials (validly cued) and on the uncued side in 20% of trials (invalidly cued) (left panel of Fig. 1). In addition, there were trials where both sides were simultaneously cued (neutral cue) and the target could appear on either side with equal probability (right panel of Fig. 1).

Trials were categorized into different conditions based on cue/target locations (attend/target inside or outside the RF), cue types (valid, neutral, or invalid), and behavioral outcomes (hit or miss). Invalidly cued trials were not analyzed due to insufficient number of trials (see ‘Data analysis’ section in Methods for more details). Since the monkeys were cued to attend to both locations in neutral blocks, we categorized those trials based on the target location instead of cue location. This resulted in the following eight conditions (four each for valid and neutral cueing): Attend-In Valid Hit (AIVH), Attend-Out Valid Hit (AOVH), Attend-In Valid Miss (AIVM), Attend-Out Valid Miss (AOVM), Target-In Neutral Hit (TINH), Target-Out Neutral Hit (TONH), Target-In Neutral Miss (TINM), Target-Out Neutral Miss (TONM)

### Attentional and Behavioral Discriminability using firing rate and LFP power

We found that differences in target onset time distribution for hits versus miss conditions had a strong effect on our results, especially at low frequencies (as discussed below in more detail). Therefore, we analyzed the data after matching the target onset time distributions of the hit and miss trials of the respective attention conditions (see Materials and Methods for details). Supplementary Figure 1A-G show the same analyses as Figure 2A-2G but before matching target onset time distribution (Supplementary Fig. 1H).

Figure 2A shows the average peri-stimulus time histogram (PSTH) across 643 electrodes across 21 sessions for the four validly cued conditions during 500 – 0 ms before the target onset (see Materials and Methods for details). Neurons fired at both the positive and negative phases of the counterphasing stimulus to produce a strong oscillatory signal at 20 Hz (De Valois et al. 1982). The average firing rates over this 500 ms interval are shown in the inset. The firing rate was higher when the monkeys attended and detected (hit) the target presented inside the RF (AIVH condition; blue) than when they attended and detected the target outside the RF (AOVH condition; red; Bonferroni corrected P = 0.01; Wilcoxon rank-sum test). This recapitulates the classic effect of visual attention on the firing rates of cortical neurons, whereby the responses to attended stimuli in the RF are enhanced, on average, relative to unattended stimuli (McAdams and Maunsell 1999). The conditions when the monkeys missed the target (AOVM; magenta and AIVM; cyan) produced a response that was intermediate between these two hit conditions since attention is expected to be on the uncued side leading to the miss.

Event related potential (ERP), obtained by averaging the LFP traces for different conditions, showed similar trends across four conditions (Fig. 2B). However, these traces showed a pronounced difference between the hit and miss conditions when the target onset time distributions were not matched (Supplementary Fig. 1B). These differences could be largely explained based on differences in target onset times for hits versus miss conditions because monkeys tended to miss the targets that appeared earlier after stimulus onset than later (Supplementary Fig. 1H). Stimulus onset produced a transient elevation in firing rate and a dip in the ERP which reached a steady state value over several hundred milliseconds (see Fig. 1A, B of Prakash et al. 2021). Thus, because our analysis window was 500 to 0 ms before the target onset (which could appear between 500-5500 ms of stimulus onset), earlier target onset times captured the transient activity for the trials in which the target appeared between 500 to 750 ms after the stimulus onset.

Figure 2C shows the change in power of each condition from AOVH condition. The effect of attention on the power spectral density (PSD) is shown by the blue trace in Figure 2C, which shows the change in power when attention is inside versus outside the RF (AIVH versus AOVH). Attention suppressed the power in lower frequencies (≤14 Hz) (Benjamini-Yekuteili FDR controlled P<0.05; Wilcoxon sign-rank test) and increased power in the gamma and high-gamma range (>36 Hz) (Benjamini-Yekuteili FDR controlled P<4.4×10^-16^; Wilcoxon sign-rank test) as well as the steady-state-visually-evoked-potential (SSVEP) frequency of 20 Hz (Prakash et al. 2021) (Benjamini-Yekuteili FDR controlled P=0.001; Wilcoxon sign-rank test). The effect of behavioral outcome (hit versus miss) was observed when attention was outside the RF and the monkey missed the target (magenta trace; AOVM versus AOVH), or by comparing the corresponding conditions for the Attend-In condition (i.e., blue versus cyan traces; AIVH versus AIVM). Upon visual inspection, low frequencies captured differences in activity related to behavioral outcome, while frequencies closer to gamma distinguished attentional state.

To better dissociate the effects of attentional location and behavioral outcome and examine how the effect of one is influenced by the other, we compared the PSDs separately by taking the difference between the attention conditions (attend-in versus attend-out) for the two behavioral outcomes: hit (Fig. 2D, green trace; same as the blue trace in Fig. 2C) and miss (Fig. 2D, yellow trace; cyan minus magenta traces in Fig. 2C). Attention-related suppression of low-frequency power (8-14 Hz) (Benjamini-Yekuteili FDR controlled P<0.01; Wilcoxon sign-rank test) and enhancement of the high-frequency power (>44Hz) (Benjamini-Yekuteili FDR controlled P<0.05; Wilcoxon sign-rank test) was also observed for the miss condition, albeit the effect was weaker than the hit condition (compare green versus yellow traces in Fig. 2D). Similarly, for behavioral effects we took the difference between the behavioral outcomes (hit – miss) for the two attention conditions: attend-in (blue trace in Fig 2E; blue minus cyan traces in Fig. 2C) and attend-out (red trace in Fig. 2E; same as the negative of the magenta trace in Fig. 2C). For the behavioral comparison (Fig. 2E), the power was different mainly in the low-frequency range for both attend-in and attend-out cases, whereas high-frequency power showed only minor differences mainly for attend-out condition.

Overall, our results show only a modest effect of behavior on LFP data that is mainly in the low-frequency range. But this is largely due to the target onset matching procedure described above. The difference in power in the very low-frequency range for the missed condition relative to the hit condition was much larger for the non-matched case (Supplementary Fig. 1E) compared to the matched case (Fig. 2E). For example, the change in 4Hz power of attend-in valid miss (AIVM) relative to attend-in valid hit (AIVH) reduced by more than three-fold in distribution matched (0.75 dB) compared to non-matched case (2.4 dB) (Fig. 2E and Supplementary Fig. 1E). This shows that the pre-target low frequency power can spuriously dominate and lead to an overestimation of its behavioral decoding performance if the differences in the distribution of target onset times between the conditions are not accounted for. For all the analyses, we therefore only used trials after matching target-onset time distributions.

Since one of the aims of the study was to compare the ability of different neural measures (such as spiking activity, LFP power, and LFP phase) to discriminate attentional conditions and behavioral outcome, we converted the LFP power and spike responses to d-prime values to compare the discriminability across neural measures (Fig. 2F and 2G; see Materials and Methods for details). Attention discriminability using firing rates (indicated by dashed lines in Fig. 2F) was higher for hit condition (green dotted line) than miss (yellow dotted line), reflecting larger separation in the means for the hits compared to miss cases (as in Fig. 2A inset). Attention discriminability with LFP power was highest in the gamma band (40 -80 Hz), was comparable to that of firing rate, and decreased slowly at higher frequencies (solid green and yellow traces in Fig. 2F; note that discriminability increases when power is averaged across frequencies, as shown below). Lower frequencies (<30 Hz) had negative d-prime (i.e., attend-in condition had lower power than attend-out as shown in Fig. 2D), but the magnitude of the discriminability was lower than that of firing rate. On the other hand, very low frequency (<14 Hz) power was best suited for discriminating the behavior (Fig. 2G) since it had the highest magnitude of discriminability, reflecting the changes in power observed in Figure 2E.

To summarize, two salient results so far are: (1) d-prime values for LFP power for attentional discriminability (Attend-In versus out; Fig. 2F) for hit condition (green trace) were highest in the gamma band, where it was comparable to d-prime of firing rate, and decreased in the high-gamma range. It was negative at low frequencies, although the magnitude was less than the gamma range. Similar trends were observed in the miss condition (yellow trace), although magnitudes were lower. (2) For behavioral discriminability (Fig. 2G) for attend-in condition (blue solid trace), d-prime values were negative at low frequencies, became positive at SSVEP frequency and in high-gamma range, comparable to the discriminability using firing rate (blue dashed line). The highest d-prime was observed for the SSVEP frequency. The effect flipped for the attend-out condition in the high-gamma range (red solid trace).

We previously showed that the single electrode LFP power in the gamma (42-78 Hz) and high-gamma (122-198 Hz) band performed better than firing rate in decoding attend-in valid hit vs. attend-out valid hit (Prakash et al. 2021). However, in our present analysis, the power in the gamma and high gamma ranges did not perform better than firing rate in discriminating the attention condition (Fig. 2F; green trace). This discrepancy is due to the variability in the spectral estimator at individual frequency bins (Jarvis and Mitra 2001), which decreases d-prime values as observed here. However, spectral variability is uncorrelated across frequencies and therefore can be reduced by summing or averaging the power over the frequencies of interest, which can improve the decoding performance (Prakash et al. 2021). We illustrate this fact by performing similar averaging by smoothening over a frequency window using Multitaper method. The extent of frequency smoothening is controlled by the time-bandwidth product (TW) which determines the number of tapers (K) to use (K = 2TW-1) for maximal spectral concentration. Increasing TW increases the frequency window over which the power is smoothed. As expected, as we increased TW, the attentional d-prime in the gamma and high-gamma band increased and exceeded the d-prime of firing rate as in our earlier study (Fig. 2H). Note that the enhanced discriminability at SSVEP was eliminated by this smoothing because the nearby frequencies did not show this discriminability. Thus, frequency smoothing improves the discriminability when the neighboring frequencies perform comparably well.

The neutral cueing condition showed similar results (compare Fig. 2A-H versus Supplementary Fig. 1I-P). The results are shown after matching the target onset times because we found similar artifacts as the valid condition when target onset times were not matched (data not shown). As in the valid condition, firing rates in neutral condition were higher when target appeared inside the RF than when it appeared outside (Fig. 2A and Supplementary Fig. 1I inset) and the ERPs were comparable across conditions (Supplementary Fig. 1J). However, there were some important differences as well. Firing rates were different for hits versus misses when target was inside the RF (blue and cyan dots in panel Supplementary Fig. 1I that correspond to TINH versus TINM conditions), but not when the target was outside the RF (TONH versus TONM). This trend was also reflected in the LFP power, with large discriminability in behavior the gamma range (which was further comparable to the firing rate discriminability) when target was in the RF (blue trace in Supplementary Fig. 1O) but not when outside (red trace in Supplementary Fig. 1O). The discriminability between target-in and target-out conditions for hit trials was highest in the gamma range and was comparable to the firing rate (green trace in Supplementary Fig. 1N) and the traces were essentially inverted for the miss condition for both firing rate and LFP power (yellow traces Supplementary Fig. 1N). Unlike the valid condition, frequency smoothening did not improve discriminability beyond firing rates (Supplementary Fig. 1P). Overall, the behavioral discriminability pattern of neutral cue was similar to the attentional discriminability of the valid condition (in other words, effect observed in Fig. 2F was similar to Supplementary Fig. 1O, while Fig. 2G was similar to Supplementary Fig. 1N).

### Discrimination of attentional state and behavioral outcome using phase

EEG studies have shown that phases at frequency bands such as alpha (8-12 Hz) are modulated by attention and are informative about behavioral performance (Thut et al. 2006; Busch et al. 2009). We therefore examined the ability of LFP phase to discriminate the attentional and behavioral states. If the LFP phases at which monkeys detect the target are consistently different from the phases at which they miss, the difference should be reflected in the ERPs, which we did not observe (Fig. 2B; although note that the presence of strong SSVEP may have affected the endogenous alpha rhythm). Because phase is a circular variable, we used the sine component of the phase vector to make it linear (like firing rate and LFP power) in order to calculate d-prime. This transformation should not affect our results since we are interested in the differences between conditions rather than their absolute values. For valid trials, the phase values were only consistent at the SSVEP frequencies (Fig. 3A), but we did not find a consistent difference across conditions at any frequency, resulting in low attentional and behavioral discriminability compared to the firing rate (Fig. 3B and 3C). Similar results were obtained when the cosine component was used instead of the sine component (data not shown).

**Figure 3:**
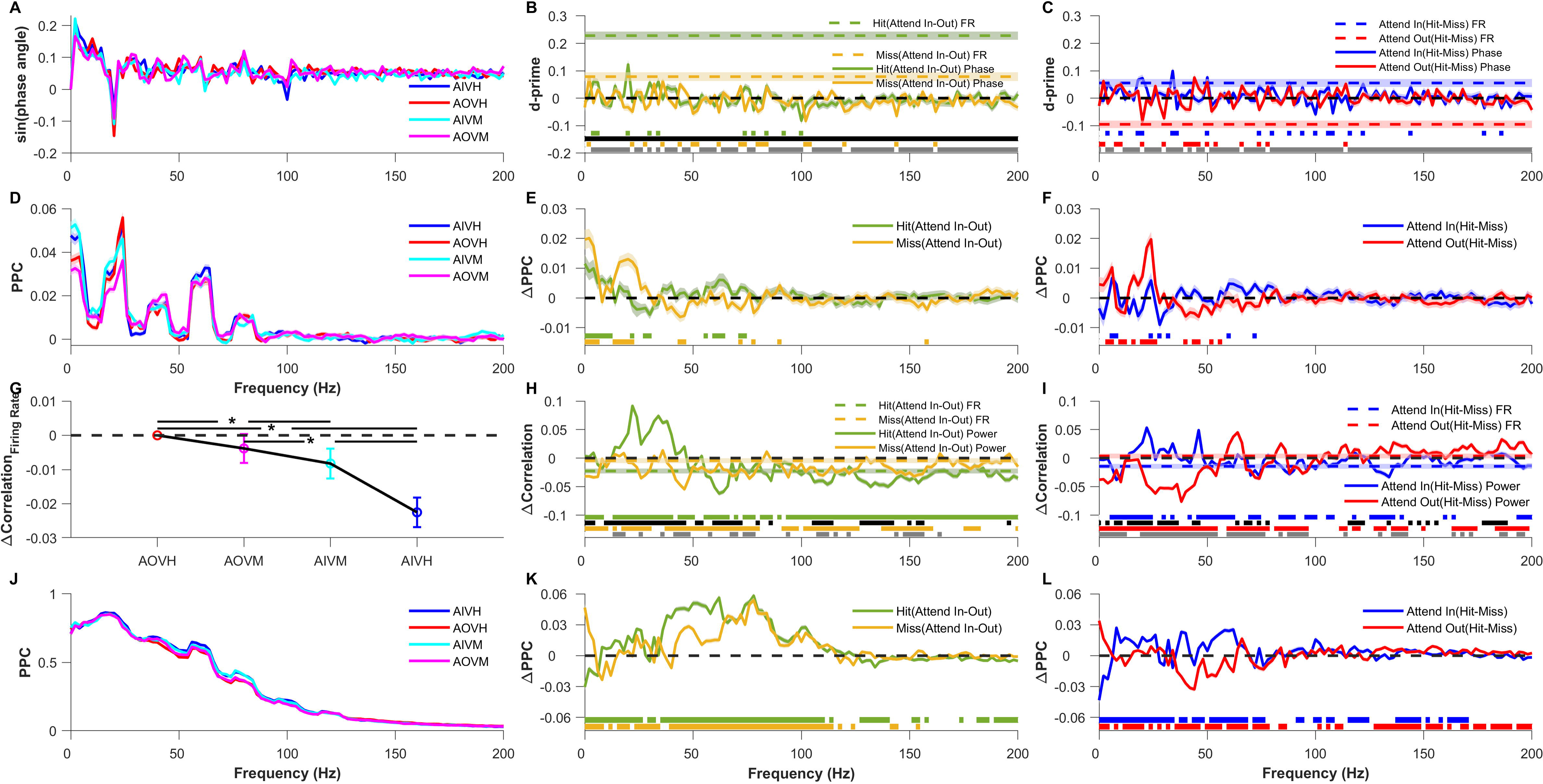
Comparison of LFP phase and pairwise phase consistency of individual electrodes and trial-wise firing rate correlation, LFP power correlation and pairwise phase consistency (PPC) of electrode pairs across validly cued attention and behavioral conditions. (A) Mean sine of the LFP phase angle for Attend-In Valid Hit (AIVH; blue), Attend-Out Valid Hit (AOVH; red), Attend-In Valid Miss (AIVM; cyan), Attend-Out Valid Miss (AOVM; magenta) conditions. (B) Mean d-prime of firing rate (dashed color lines) and sine of the LFP phase angle (solid lines) between attend-in and attend-out conditions for hit (green) and miss (yellow) conditions. The horizontal dashed black line marks the zero of the y-axis. Horizontal green and yellow patches at the bottom indicate the frequencies at which the d-prime of LFP phase angle for hit and miss condition respectively are significantly greater than zero (Wilcoxon signed-rank test, Benjamini-Yekutieli FDR controlled p<0.05 under unknown dependency). Horizontal black and grey patches at the bottom indicate the frequencies at which the d-prime of LFP phase angle is significantly different from d-prime of firing rate for hit and miss condition respectively (Wilcoxon rank-sum test, Benjamini-Yekutieli FDR controlled p<0.05 under unknown dependency). Negative d-prime values were multiplied by -1 before performing the significance test. (C) Mean d-prime of firing rate (dashed color lines) and sine of the LFP phase angle (solid lines) between hit and miss conditions for attend-in (blue) and attend-out (red) conditions. The horizontal dashed black line marks the zero of the y-axis. Horizontal blue and red patches at the bottom indicate the frequencies at which the d-prime of LFP phase angle for attend-in and attend-out condition respectively are significantly greater than zero (Wilcoxon signed-rank test, Benjamini-Yekutieli FDR controlled p<0.05 under unknown dependency). Horizontal grey patches at the bottom indicate the frequencies at which the d-prime of LFP phase angle is significantly different from d-prime of firing rate for attend-out condition. (Wilcoxon rank-sum test, Benjamini-Yekutieli FDR controlled p<0.05 under unknown dependency). Note that difference between d-prime for LFP phase angle and firing rate for attend-in condition was not significant at any frequency (Wilcoxon rank-sum test, Benjamini-Yekutieli procedure for FDR control under unknown dependency), hence no patch is shown for that comparison. Negative d-prime values were multiplied by -1 before performing the significance test. (D) Mean single electrode pairwise phase consistency (PPC; see Materials and Methods for details) for the four conditions as in (A). (E) Mean change in single electrode PPC between attend-in and attend-out conditions for hit (green) and miss condition (yellow). The horizontal dashed black line marks the zero of the y-axis. Horizontal green and yellow patches at the bottom indicate the frequencies at which the change in PPC for hit and miss condition respectively are significantly greater than zero (Wilcoxon signed-rank test, Benjamini-Yekutieli FDR controlled p<0.05 under unknown dependency). (F) Mean change in single electrode PPC between hit and miss conditions for attend-in (blue) and attend-out condition (red). The horizontal dashed black line marks the zero of the y-axis. Horizontal blue and red patches at the bottom indicate the frequencies at which the change in PPC for attend-in and attend-out condition respectively are significantly greater than zero (Wilcoxon signed-rank test, Benjamini-Yekutieli FDR controlled p<0.05 under unknown dependency). In (A) – (F) mean and s.e.m are computed like in Fig. 2A – 2G. (G) Mean change in trial-wise firing rate correlations for the four conditions relative to Attend-Out Valid Hit (AOVH) condition. Dashed black line indicates the zero of y-axis. Asterisks indicate the conditions between which the differences in trial-wise correlation are significant (Wilcoxon rank sum test; Bonferroni corrected p<0.05) between the conditions. (H) Mean change in trial-wise LFP correlations (solid lines) and firing rate correlation (dashed color lines) between attend-in condition and attend-out condition for hit (green) and miss trials (yellow). The dashed black line indicates the zero of y-axis. (I) Mean change in trial-wise LFP correlation (solid lines) and firing rate correlation (dashed color lines) between hit condition and miss condition for validly cued attend-in (blue) and attend-out (red) conditions. The dashed black line indicates the zero of y-axis. (J) Mean pairwise phase consistency (PPC) across trials for the four conditions as in (A). (K-L) Same as (E-F) but for pairwise phase consistency (PPC) across trials. In (G) – (L) Mean was taken across 50 bootstrap samples of mean across 5754 pairs and shaded lines and error bar indicate the bootstrap mean of s.e.m across 5754 electrode pairs. In (H), (I), (K) and (L), significance lines shown in the bottom are calculated using the same approach as (B), (C), (E) and (F).

We previously showed that pairwise phase consistency (PPC) across pairs of electrodes, which reflects the consistency of the phase difference between electrodes across trials, varied with attentional condition (see Fig. 2. of Prakash et al. 2021). We next tested whether absolute phase of a single electrode also showed any consistency across trials. For this analysis, we computed the PPC, which is an unbiased estimator of the square of the coherence (Vinck et al. 2010), but replaced the phase difference between a pair of electrodes with the absolute phase of one electrode for each trial. This single-electrode PPC is expected to be high if the absolute phase is consistent across trials. However, PPC was high only at the SSVEP frequency (the peak is seen at 24 Hz instead of 20 due to smoothening effect of Multitaper) and its harmonics (Fig. 3D). No consistent trends were observed when we compared across conditions (Fig. 3E and 3F). Note that since PPC is computed across trials, it is not a single-trial measure and therefore we cannot obtain d-prime values like above. In summary, the absolute phase of the LFP was much less informative about the attentional location or behavioral state than firing rates, and no difference in the consistency of the phase across trials was observed across conditions. Similar results were obtained for the neutral cue (Supplementary Fig. 2A-F).

### Effect of attention and behavioral outcome on pairwise measures

Conventional measures based on pairs of electrodes, such as correlation and phase consistency, are computed across trials. Before exploring single-trial estimates of such measures, which are required to estimate discriminability, we evaluated the effects of attention and behavior the traditional way by computing these measures across trials. Consistent with previous studies (Cohen and Maunsell 2009; Mitchell et al. 2009; Mayo and Maunsell 2016), attention reduced correlations in firing rate across pairs of electrodes (AOVH versus AIVH; Fig. 3G). The correlation for missed conditions were intermediate to the hit condition (AOVM and AIVM; Fig 3G), consistent with the changes in firing rate as shown in Figure 2A inset. Correlations between LFP power recorded from two electrodes increased with attention at SSVEP frequency (20 Hz) and its harmonic but reduced for frequencies higher than 50 Hz for valid hit condition (Fig. 3H; green trace). Such modulations were not observed when the animals missed the trials (yellow trace in Fig. 3H). Differences between behavioral outcomes (hit versus miss; Fig. 3I) were small in the high-gamma range, mirroring the small changes observed in firing rate correlations as well. Interestingly, lower frequencies including the SSVEP frequency showed larger differences for Attend-out condition (red trace in Fig. 3I) which is similar to the strong effect of behavioral differences on SSVEP power observed earlier (Fig. 2G). Overall, these results are consistent with previous studies that showed a reduction in spike count correlation (Cohen and Maunsell 2009; Mitchell et al. 2009) and LFP power correlation except at SSVEP (Prakash et al. 2021).

Next, we analyzed the PPC between the LFP signals from pairs of electrodes (Fig. 3J-L). PPC increased for valid condition when attention was cued into the RF over a broad range of frequencies between 36 Hz to 106 Hz (Fig. 3K; green trace), although we did not observe much difference beyond 120 Hz even though LFP power was elevated in that range (compare with the green trace in Fig. 2D). Reasons for this difference between high-frequency power and PPC are discussed elsewhere (Ray 2022). Interestingly, PPC was more sensitive to attentional differences (Fig. 3K) than behavioral differences (Fig. 3L).

For neutral condition (Supplementary Fig. 2G-L), the effect of target location (Supplementary Fig. 2H and Supplementary Fig. 2K) was somewhat similar to behavioral effect of the valid condition (Fig. 3I and Fig. 3L) and the behavioral effect (Supplementary Fig. 2I and Supplementary Fig. 2L) was similar to attentional effect of the valid condition (Fig. 3H and Fig. 3K), as observed previously with LFP power (Supplementary Fig. 1N was similar to Fig. 2G with the two traces flipped horizontally, while Supplementary Fig. 1O was similar to Fig. 2F with one trace showing salient effect and the other one close to zero).

### Discrimination of attentional state and behavioral outcome using pairwise measures

Modulation of pairwise measures by attentional state and behavioral outcome motivated us to develop methods to obtain the same measures in single trials, which could then be used to obtain discriminability values (d-prime) like the power and phase measures we used earlier. We devised two independent methods to estimate correlation and phase consistency within a single trial. In the first method, we divided the spiking and LFP signal of a single trial into ten non-overlapping bins of 50 ms each and computed firing rate, LFP power, and phase in each bin. We then calculated the correlation and LFP-LFP phase consistency between electrode pairs across bins (Fig.4, left panel). In the second approach we used the Multitaper method where a single trial LFP signal was multiplied by nine orthogonal Slepian tapers to obtain nine independent estimates of LFP power and phase (Fig. 4, right panel) and computed their correlation and phase consistency (see Materials and Methods for more details). The first method is more direct because power and LFP values are obtained from non-overlapping segments of data, although the frequency resolution (reciprocal of the analysis duration in seconds) becomes 20 Hz. The second method uses the same length of data (500 ms) to get several estimates of power and phase (which are independent because the tapers are orthogonal) and therefore has a frequency resolution of 2 Hz. We also used a method based on Hilbert transform to estimate instantaneous power and phase at each time point within a trial and then computed correlation and phase consistency across time (Spyropoulos et al. 2024).

**Figure 4:**
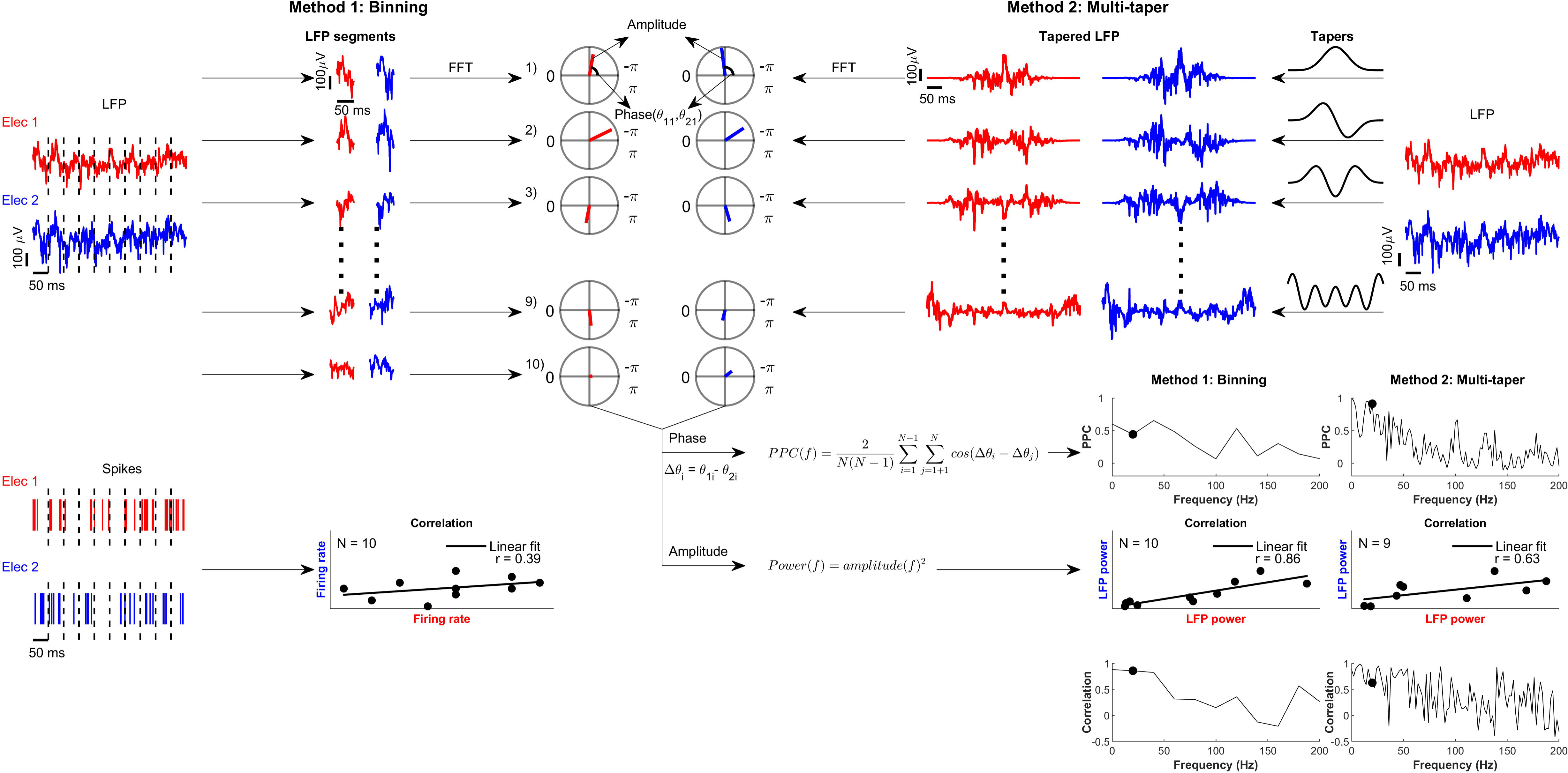
Illustration of methods used for estimating correlation and pairwise phase consistency within a trial. Binning method (left panel): A single-trial LFP signal of 500 ms duration from two electrodes is first divided into 10 non-overlapping bins of 50 ms each. Then, the power and phase of each 50 ms LFP segments of the two electrodes were estimated by Fourier transform. Subsequently, we estimated the power correlation and pairwise phase consistency across the ten bins between the pair of electrodes at each frequency between 0-200 Hz with 20 Hz resolution. Similarly, the spikes from the two electrodes were also binned and firing rate correlations between the two electrodes were computed across bins (lower left panel). These correlations were subsequently shuffle corrected (see methods). Multitaper method (right panel): We used an alternate method to obtain multiple estimate of power and phase using the Multitaper method. LFP signal of 500 ms duration of two electrodes were first multiplied by nine orthogonal Slepian tapers of same length, then power and phase of these nine tapered LFP signals were obtained by Fourier transform. Power correlation and PPC were calculated between electrode pairs across tapers at frequencies between 0-200 Hz with 2Hz resolution. The black dot in the PPC vs frequency and correlation vs frequency plots indicate the frequency whose amplitude and phase are shown in the polar plot in top panel and whose power scatter plot is shown.

Firing rate correlations were reduced with attention for hit trials even when computed over a single trial (Fig. 5A, dashed green line), but not reduced in miss trials (Fig. 5A, dashed yellow line), consistent with the across-trial correlation measures (Fig. 3H). Even the single-trial LFP power correlation computed using the binning method showed similar effects as that of the traditional correlation measure (compare solid green and yellow traces in Fig. 5A with the same traces in Fig. 3H). Because these were single-trial measures, we could now calculate and compare the discriminability by calculating d-prime values in the same way as before (Fig. 5B). LFP correlations in the high-gamma range could distinguish the attention conditions slightly better than firing rate correlation when the targets were correctly detected (Fig. 5B; green trace), although these d-prime values were considerably lower than the ones obtained using firing rate/LFP power (compare with similar traces in Fig. 2F). Similar trends were obtained for behavioral comparison: (1) single trial correlations (Fig. 5C) mirrored the results obtained using across-trial correlations (Fig. 3I), (2) magnitude of discriminability (Fig. 5D) was higher in the high-gamma range where it was comparable to firing rate correlations for the attend-in condition (blue traces in Fig. 5D), and (3) for the attend-out condition the low-gamma frequencies were more discriminable (red trace in Fig. 5D) than firing rate for that condition. In summary, we found that our single trial estimates of correlation showed modulations similar to the traditional trial-wise correlation and the discriminability of high-gamma and firing rate correlations were comparable.

**Figure 5:**
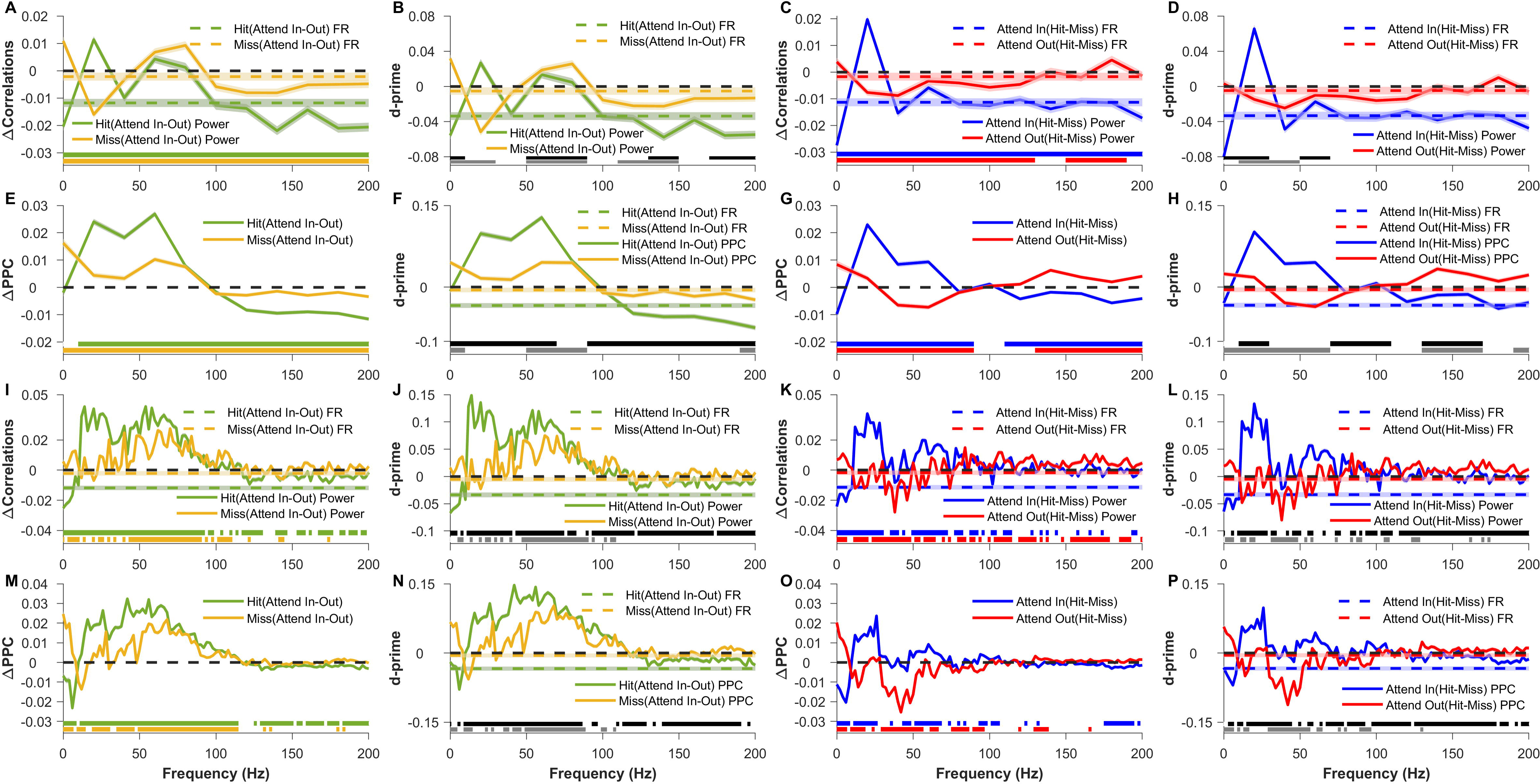
Comparison of single trial bin-wise firing rate correlation, bin-wise and taper-wise LFP power correlation and pairwise phase consistency (PPC) across validly cued attention and behavioral conditions. (A) Mean change in single trial bin-wise LFP power correlation (solid lines) and firing rate correlation (dashed color lines) between validly cued attend-in and attend-out conditions for hit (green) and miss (yellow) conditions. Single trial bin-wise correlation was measured by dividing the analysis period of 500 ms into 10 non-overlapping bins and measuring the firing rate or power correlation across bins. Since power was estimated for signals of 50 ms duration, the frequency resolution is 20 Hz. Dashed black line indicates the zero of y-axis. (B) Mean d-prime of bin-wise LFP power correlation (solid lines) and firing rate correlation (dashed color lines) between validly cued attend-in and attend-out conditions for hit (green) and miss (yellow) conditions. Dashed black line indicates the zero of y-axis. (C) Mean change in bin-wise LFP power correlation (solid lines) and firing rate correlation (dashed color lines) between hit and miss conditions for validly cued attend-in (blue) and attend-out (red) conditions. Dashed black line indicates the zero of y-axis. (D) Mean d-prime of bin-wise LFP power correlation (solid lines) and firing rate correlation (dashed color lines) between hit and miss condition for validly cued attend-in (blue) and attend-out (red) conditions. Dashed black line indicates the zero of y-axis. (E) – (H) Same as (A) – (D) but for single trial bin-wise PPC. Single trial bin-wise PPC was computed across the same 10 non-overlapping bins used to compute power correlations. In (A) – (H) the colored patches at the bottom indicate the significance like that in figure 3 but here the FDR was controlled using Benjamini – Hochberg procedure. (I) – (P) Same as (A) – (H) but for single trial taper-wise power correlation and PPC. Here power correlation and PPC are computed by using multi-taper method. Specifically, we use TW=5 to get 9 (2TW-1) estimates of power and phase values per trial and compute the power correlation or PPC between electrode pairs across these nine values. Unlike (A) – (H) FDR was controlled by using Benjamini-Yekutieli procedure under unknown dependency. In (A) – (P) Mean was taken across 50 bootstrap samples of mean across 5754 pairs and shaded lines indicate the bootstrap mean of s.e.m across 5754 electrode pairs.

Results of single-trial bin-wise PPC (Fig. 5E and 5G) also showed similar trends as that of conventional trialwise PPC (Fig. 3K and 3L). However, the magnitude of d-prime values was considerably higher (Fig. 5F and 5H) compared to that of correlations in firing rate/power, suggesting that single trial PPC, especially in the gamma range, was a useful measure to discriminate attention or the behavioral outcome.

Although bin-wise PPC reflected most of the trends seen in trialwise PPC, the binning method was inherently limited in its frequency resolution. To overcome this limitation, we used the Multitaper method and computed correlation and PPC across tapers (Method 2 in Fig. 4). The binning and Multitaper methods produced similar results, validating our approaches. For correlations (Fig. 5I-L), while the trends were generally similar to the bin-wise approach (compare with Fig. 5A-D), we found two important differences. First, while the high-gamma band beyond 100 Hz essentially reflected the trends observed in firing rates for the bin-wise method (e.g., Fig. 5A), the Multitaper method produced negligible correlation differences above 100 Hz (e.g., Fig. 5I). On the other hand, the difference in LFP correlations between the conditions improved remarkably in the gamma band and below (<60 Hz) and hence the discriminability index increased threefold compared to the binning method (compare Fig. 5B and 5D with Fig. 5J and 5L; note the difference in scales). In contrast, the PPC measure (Fig. 5M-P) remained comparable across both methods (Fig. 5E-H). Among the electrode pair measures, PPC in the low to mid frequency range was the most useful measure to discriminate attention and behavior. For the neutral conditions (Supplementary Fig. 3), the trends were similar to their across-trial counterparts (Supplementary Fig. 2) and reflected the same results as observed for valid trials. Single-trial estimates of power correlation and phase consistency calculated using Hilbert transform method (Supplementary Fig. 4) yielded largely comparable results.

Finally, because our analyses in Figure 2H demonstrated that discriminability differs as a function of smoothening over frequency ranges, we computed all the LFP measures according to frequency bands of alpha (8-12 Hz), SSVEP (18-22 Hz), gamma (40-80 Hz), high-gamma (120-200 Hz) and summarized our results in Figure 6. Same results for the neutral condition are shown in Supplementary Figure 5. Our key findings are summarized as follows: 1) LFP power in the high-gamma and gamma bands were best suited for discriminating “attend-in” vs. “attend-out” attentional state (d-prime values of 0.32 ± 0.01 and 0.3 ± 0.01; Fig. 6A). Firing rate was the next most informative measure (d-prime of 0.23 ± 0.01; Fig. 6A), but discriminability using gamma was significantly greater than that of firing rate (p = 4.6×10^-5^; Wilcoxon rank sum test). Alpha power also had strong discriminability (d-prime of -0.14 ± 0.01), although the magnitude was significantly less than firing rate (p = 1.1×10^-6^; Wilcoxon rank sum test). 2) While gamma, high-gamma, and spiking activity could all be reliably used to discriminate attentional state, alpha power, SSVEP power, and SSVEP phase were the most informative measures for discriminating behavioral outcome (Fig. 6B). In general, firing rates and high-frequency LFP (gamma and high-gamma) were not very useful to discriminate behavior. 3) Among the phase-based measures, single trial PPC in the gamma band was the most informative of the attentional state, especially when calculated using the Multitaper method (d-prime of 0.23 ± 0.004; purple bar in Fig. 6A), on par with the firing rate (p = 0.95; Wilcoxon rank sum test). 4) The performance of the neural measures in the discriminating behavioral outcome for neutral condition (Supplementary Fig. 5B) was similar to the discrimination of attentional conditions for valid condition (Fig. 6A).

**Figure 6:**
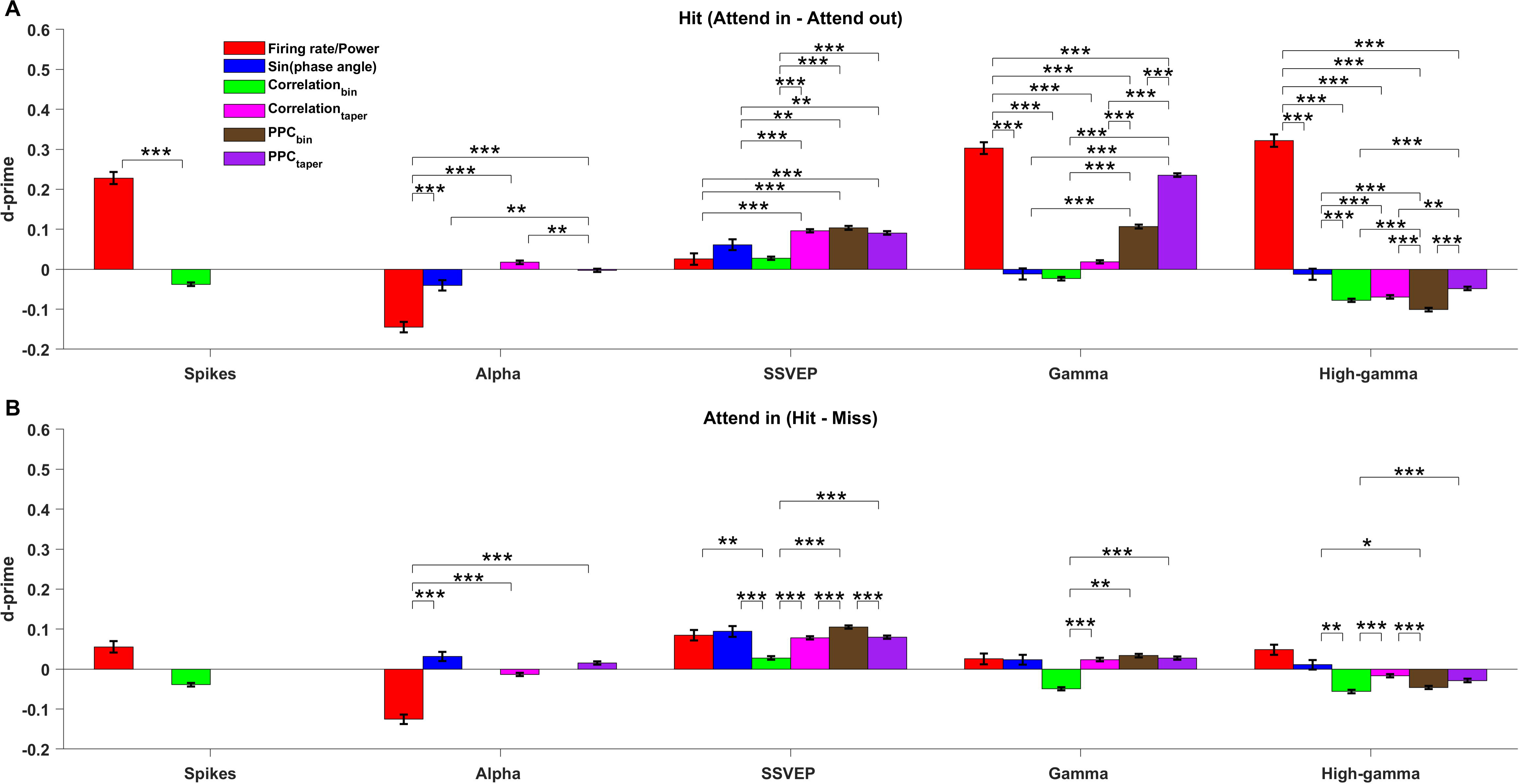
Summary plot comparing across the d-primes of all the measures for spikes, LFP power and phase in different frequency bands for validly cued attention and behavioral comparison. (A) D-prime of firing rate/ LFP power (red), sine of the phase angle (blue), firing rate/ LFP power correlation across the bins (green), LFP power correlation across the tapers (magenta), pairwise phase consistency (PPC) across bins (brown), PPC across tapers (purple) for spikes, alpha (8-12 Hz), SSVEP (18-22 Hz), gamma (40-80 Hz) and high-gamma (120-200 Hz) power between attend-in and attend-out conditions for hit trials. Note that for spikes, only d-primes of firing rate and bin-wise correlation exist because rest of the measures cannot be computed for the spike data whereas for alpha-band, d-prime of bin-wise measures (CorrelationBin and PPCBin) are absent because the frequency resolution in the binning method was 20 Hz. ‘***’ indicates p-value<0.001, ‘**’ indicates 0.001<p-values<0.01, ‘*’ indicates the 0.01<p-values<0.05 and no asterisks are shown for p-values>0.05. Negative d-prime values were converted to positive values by multiplying with -1 before performing the significance test since we were comparing only the magnitude. All the p-values are Bonferroni corrected for multiple comparisons. (B) Same as (A) but for the d-prime between the hit and miss conditions of Attend-in condition.

## Discussion

We studied the effect of attentional (cue inside vs outside RF) location and behavioral outcome (hit vs miss) on distinct frequency bands of the LFP. Our analyses illustrated potential issues caused by visual transients that can differentially affect measures of behavioral outcome, which could be corrected by matching the target onset distributions for different conditions. While high-frequency LFP power in the gamma (42-78 Hz) and high-gamma (122-198 Hz) band decoded the attentional state best, low frequency power in the alpha (8-12 Hz) and SSVEP (18-22 Hz) range decoded the behavior well. We also used single-trial estimates of correlation and pairwise phase consistency (PPC) which showed similar attention and behavioral modulations as their trial-wise counterparts. Although these metrics did not produce the best decoding performance in our dataset, but they will likely be useful for future investigations of the dynamics of attention. Phase-based measures performed poorly compared to power-based measures. Nonetheless, among the phase-based measures, taper-wise PPC in the gamma band was the best, performing on par with the firing rate in decoding attention.

### Comparison with previous studies

LFP high-gamma power has been shown to reflect the average firing rate of the neuronal population around the microelectrode tip (Ray et al. 2008; Ray and Maunsell 2011). Unlike LFP, the increase in high-gamma power in electrocorticogram (ECoG) – a signal which integrates over a larger population than LFP – cannot be accounted solely by increases in firing rate and has been hypothesized to reflect more synchronous or correlated activity in the neural population (Ray et al. 2008). Interestingly, we found that attentional discriminability of LFP high-gamma power was higher than that of firing rate which suggests that, like ECoG, high-gamma power of LFP could also partially reflect firing rate correlations and synchrony (in addition to the average firing rate) which also get modulated with attention (Cohen and Maunsell 2009). In addition, improved performance of higher LFP frequencies could simply be due to optimal averaging of responses in the LFP signal over the spatial scale over which attention operates, as argued previously (Prakash et al. 2022).

Our results are consistent with the findings of human EEG studies (Thut et al. 2006; Hanslmayr et al. 2007; Busch et al. 2009) which showed that alpha power was lower when subjects detected a visual target than when they missed it, although we note that the endogenous alpha rhythm in our study could be compromised because of the strong 10 Hz counterphasing stimulus. However, care should be taken to match the distribution of target onset times for hit and miss trials while analyzing the data in the pre-target period to avoid the effect of stimulus onset related transients that can artificially inflate the behavioral effects on low frequency power, especially in a change detection task which is often used in attention studies.

We found the performance of phase-based measures underwhelming given many studies have shown the importance of phase in attention and behavioral performance (Busch and VanRullen 2010; Fiebelkorn et al. 2018; Rohenkohl et al. 2018). For example, attention improved the consistency of gamma-band phase relations between two neuronal groups, which is thought to be crucial for the effective relay of information (Fries et al. 2001). Attention is also suggested to operate in a rhythmic manner which is mediated by the phase of alpha and theta band (Busch and VanRullen 2010; Landau and Fries 2012; Fiebelkorn et al. 2018), although a recent study has questioned these results (Brookshire 2022). Previous LFP studies which used reaction time as the behavioral readout have shown that absolute phase (Ni et al. 2016) and phase coherency (Womelsdorf et al. 2006) in the gamma band to be predictive of behavior. Despite these mechanistic implications of phase on attention, we did not find any effect of absolute phase. Even with pairwise measures, only gamma-band PPC managed to perform well in discriminating attentional state, although not as well as power-based measures. Interestingly, the attentional discriminability of gamma-band PPC was comparable to that of firing rate, contrary to a recent finding where V4 firing rate decoded attention better than inter-areal gamma-band PPC between V1 and V4 (Spyropoulos et al. 2024). Furthermore, several EEG studies (Busch and VanRullen 2010; Harris et al. 2018), showed that the detection performance at the cued location systematically depended on the phase of the alpha-band, their performance in discriminating the behavioral outcome was marginal. The performance of SSVEP phase measures, on the other hand, was comparable to that of its power. Our results of LFP power-based measure performing better than phase-based measures are consistent with a similar finding in epidural ECoG recordings in macaque visual area V1 and V4 (Rotermund et al. 2013).

In our study, the absolute phase of LFP did not perform well in discriminating attention or behavioral outcome, possibly because the phase values were estimated over a coarser timescale of 500 ms, during which phase values at higher frequencies were repeated multiple times. High-frequency oscillations often occur in short bursts (Feingold et al. 2015; Lundqvist et al. 2016); for example, gamma oscillations occur in bursts with a duration of 100-300 ms (Burns et al. 2011; Xing et al. 2012; Chandran KS et al. 2018), such that a single phase value estimated over 500 ms may not capture the changes occurring over shorter timescales. We minimized this limitation by calculating phase over a shorter time-period of 200 ms before the target onset, but results did not change appreciably (data not shown). Further, earlier studies (Busch et al. 2009; Ni et al. 2016; Harris et al. 2018) have compared absolute phases for different behavioral measures using different techniques. For example, in previous studies (Busch et al. 2009; Harris et al. 2018), inter-trial coherence (ITC) – a measure that quantifies the concentration of phase vectors across trials – was computed separately for trials of hit and miss conditions. These ITC values were then compared against the overall ITC computed across all the trials from both conditions combined to quantify the differences in phase concentration as well as the average phase value between the two conditions. This is essentially a trial-averaged measure, which is different from using the sine or cosine of the phase angle of individual trials as done in this study since our main goal was to compute d-prime values for all measures for a common comparison, which cannot be computed for previous methods.

### Shared and distinct effects of attention and behavior on LFP

Although subjects are better at detecting the target when it appears at the cued location, they occasionally miss the target. If this is due to the misallocation of attention, the modulation of neural activity by attention (attend-in vs attend-out) and behavioral outcome (hit vs miss) should be similar, which was not the case. One reason could be that in a missed trial, the overall attentional (or arousal) level could be low (rather than focused on the incorrect side), which would further be reflected in the power of low-frequency rhythms such as alpha (Hong et al. 2014; van Kempen et al. 2019; Johnston et al. 2022). Further, a behavioral report involves a decision variable, and hence relates to different aspects such as bias or criterion, as compared to changes in sensitivity (Sridharan et al. 2014; Banerjee et al. 2019; Luo and Maunsell 2019). Some studies have suggested that bias could be reflected in alpha power (Limbach and Corballis 2016; Iemi et al. 2017; Samaha et al. 2020; but see Podvalny et al. 2021 and Zhou et al. 2021) potentially leading to better discriminability of behavioral outcome using alpha power. Saccade preparation is another factor which can modulate the low-frequency oscillations in V4 (Steinmetz and Moore 2014) that might contribute to better behavioral discriminability. However, a recent study which used the same experimental data found no difference in the microsaccade direction – a potential indicator of saccade preparation (Hafed and Krauzlis 2012) - between hits and miss trials (Willett and Mayo 2023).

Our results also relate to potential roles of fast (gamma) and slow (alpha/beta) oscillations in mediating feedforward and feedback communication across brain areas, as proposed by some studies (van Kerkoerle et al. 2014; Bastos et al. 2015). This notion implies that attentional modulation somehow affects the feedforward pathway while behavioral outcome (target detection) affects feedback, which is unlikely for several reasons. First, attention is shown to enhance gamma-band causal influence (Granger causality) in both feedforward and feedback directions between area V4 and V1 and within V4 (Ferro et al. 2021). Second, attention signals are shown to originate from higher areas like frontal eye field (FEF) which sends a long-range feedback connections to V4 (Gregoriou et al. 2009). On the other hand, the behavioral outcome effects could be mediated through the feedback pathway because an enhancement of alpha band activity was observed in lateral intraparietal area (LIP) when the perceptual sensitivity was low which can lead to a miss and a boost in the beta band activity in FEF when the perceptual sensitivity was high which can lead to a hit (Fiebelkorn et al. 2018). The inter-areal coherence usually increases in the frequency bands which have higher power (Schneider et al. 2021) and consequently the behavioral outcome information can propagate through the low-frequency mediated feedback signal to lower areas like V4. Therefore, our results show that modulation of high or low frequency oscillations might not always result from the exclusive engagement of feedforward or feedback pathways respectively, but instead can arise from both feedforward and feedback, at least for attentional effects.

### Decoding performance in the neutral condition

There were similarities as well as stark differences in discriminability of attention and behavioral outcomes in neutral condition. Like in valid condition, LFP power was better at decoding than phase measures. However, high-frequency oscillations were better in decoding behavioral outcome than attention location, in contrast to valid condition. The difference in the discriminability pattern could be due to the differences in the strategies employed in the two cue conditions. In the neutral condition, subjects likely had to divide their attention or switch between the two locations since the target was equally likely to appear at either location. The similarity between the decoding pattern of behavioral outcome (Hit vs Miss) in neutral condition and attention (attend-in vs attend-out) in valid condition suggests that the focus of attention was outside the receptive field whenever the subjects missed the target that appeared inside the receptive field, since the target was equally likely to occur on either side, making it similar to the attend-in and attend-out of the valid condition. This explanation holds only if the subjects were switching between the two locations. However, we do not know the exact strategy the subjects used during neutral cue; future experiments that control for the strategies employed by the subject may help resolve the similarities observed between the decoding patterns of different states of the valid and neutral conditions.

Overall, the differential ability of distinct LFP bands in discriminating attention and behavioral states not only highlights the usefulness of LFP in segregating the information related to different cognitive and behavioral state but also provides insights into their underlying mechanisms.

## Acknowledgements

We thank Dr. John Maunsell for his help in experimental design and data collection.

## Author contributions

Conceptualization, S.S.P., J.P.M. and S.R.; Methodology, S.S.P., J.P.M. and S.R.; Formal Analysis, S.S.P. and S.R.., Investigation, J.P.M.; Writing – Original Draft, S.S.P.; Writing – Review & Editing, S.S.P., J.P.M. and S.R.; Funding Acquisition, J.P.M. and S.R.; Resources, J.P.M. and S.R.; Supervision, J.P.M. and S.R.

## Data and Code Availability

All the data and original code required to replicate the results reported in this paper are accessible from the Github repository: https://github.com/supratimray/MayoProject2

## Figure legends

**Supplementary Figure 1:**
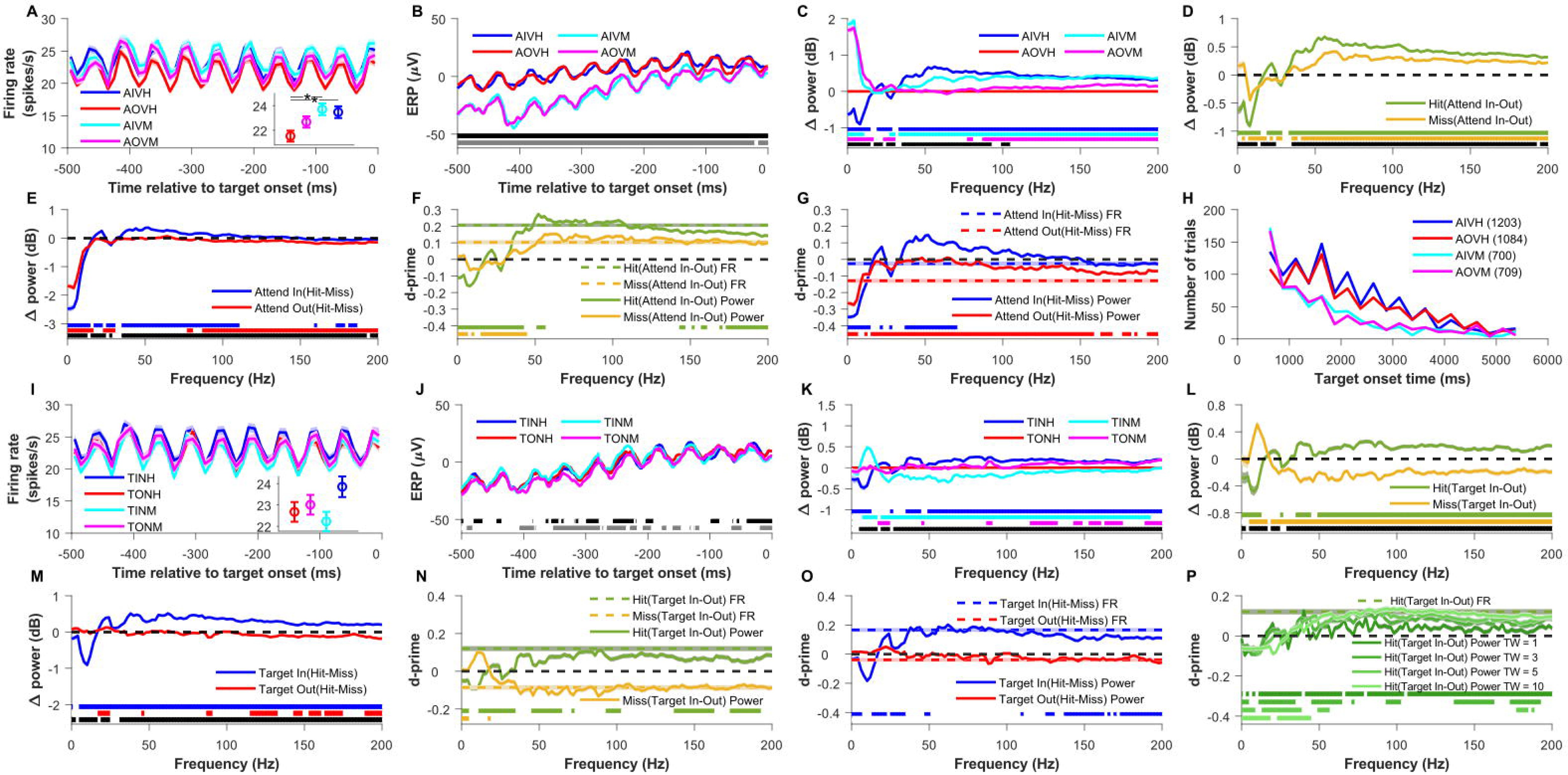
Comparison of firing rate (FR) and local field potential (LFP) power across (1) validly cued attention and behavioral conditions for non-matched target onset time distributions and (2) neutrally cued attention and behavioral conditions for matched target onset time distributions. (A) – (G) Same as Figure 2 (A) – (G) but for the case where the target onset time distributions of hit and miss conditions were not matched. Here the mean is taken across 677 electrodes recorded across 22 sessions in two monkeys. Shaded lines and error bars (not visible for most traces) indicate the s.e.m across the 677 electrodes. (H) Frequency distribution of target onset time for the four validly cued conditions of all the sessions. The number in the brackets indicate total number of trials in the respective conditions. (I) Mean peri-stimulus time histogram (PSTH) relative to the target onset time for the neutrally cued conditions in which attention was cued to both visual hemifields simultaneously and target could appear at either of the location with 50% probability. Unlike the valid cue condition where the conditions were divided based on attention location, here the conditions are divided based on where the target eventually appeared, namely Target-In Neutral Hit (TINH; blue), Target-Out Neutral Hit (TONH; red), Target-In Neutral Miss (TINM; cyan), Target-Out Neutral Miss (TONM; magenta). Inset shows the mean firing rate over the same time period as PSTH for the four conditions. Mean is first taken across 659 electrodes recorded across 21 sessions in two monkeys and then averaged across 50 bootstrap iterations. Shaded lines and error bars indicate the bootstrap mean of s.e.m across the 659 electrodes. (J) Mean ERP for the four conditions described in A. (K) Mean change in power spectral density for all the neutrally cued conditions relative to Target-Out Neutral Hit (TONH) condition. (L) Mean change in LFP power spectral density in decibels between Target-In and Target-Out conditions for hit (TINH and TONH; green) and miss (TINM and TONM; yellow). The horizontal dashed black line marks the zero of the y-axis. (M) Mean change in LFP power spectral density between hit and miss condition for neutrally cued Target-In (TINH and TINM; blue) and Target-Out (TONH and TONM; red) conditions. The horizontal dashed black line indicates the zero of the y-axis. (N) Mean d-prime of LFP power (solid lines) and firing rate (dashed color lines) between neutrally cued attend-in and attend-out conditions for hit (green) and miss (yellow) conditions. The horizontal dashed black line marks the zero of the y-axis. (O) Mean d-prime of LFP power (solid lines) and firing rate (dashed color lines) between hit and miss condition for neutrally cued Target-In (blue) and Target-out (red) conditions. The horizontal dashed black line indicates the zero of the y-axis. (P) Mean d-prime of LFP power between Target-In and Target-Out conditions for hit condition where LFP power is estimated by Multitaper method using time-frequency bandwidth product (TW) of 1 (darkest green), 3 (dark green), 5 (light green) and 10 (lightest green). The dashed green line indicates the d-prime of firing rate between the same conditions. The horizontal dashed black line marks the zero of the y-axis. Asterisks and horizontal color patches at the bottom of each panel indicate the significance level like in Figure 2.

**Supplementary Figure 2:**
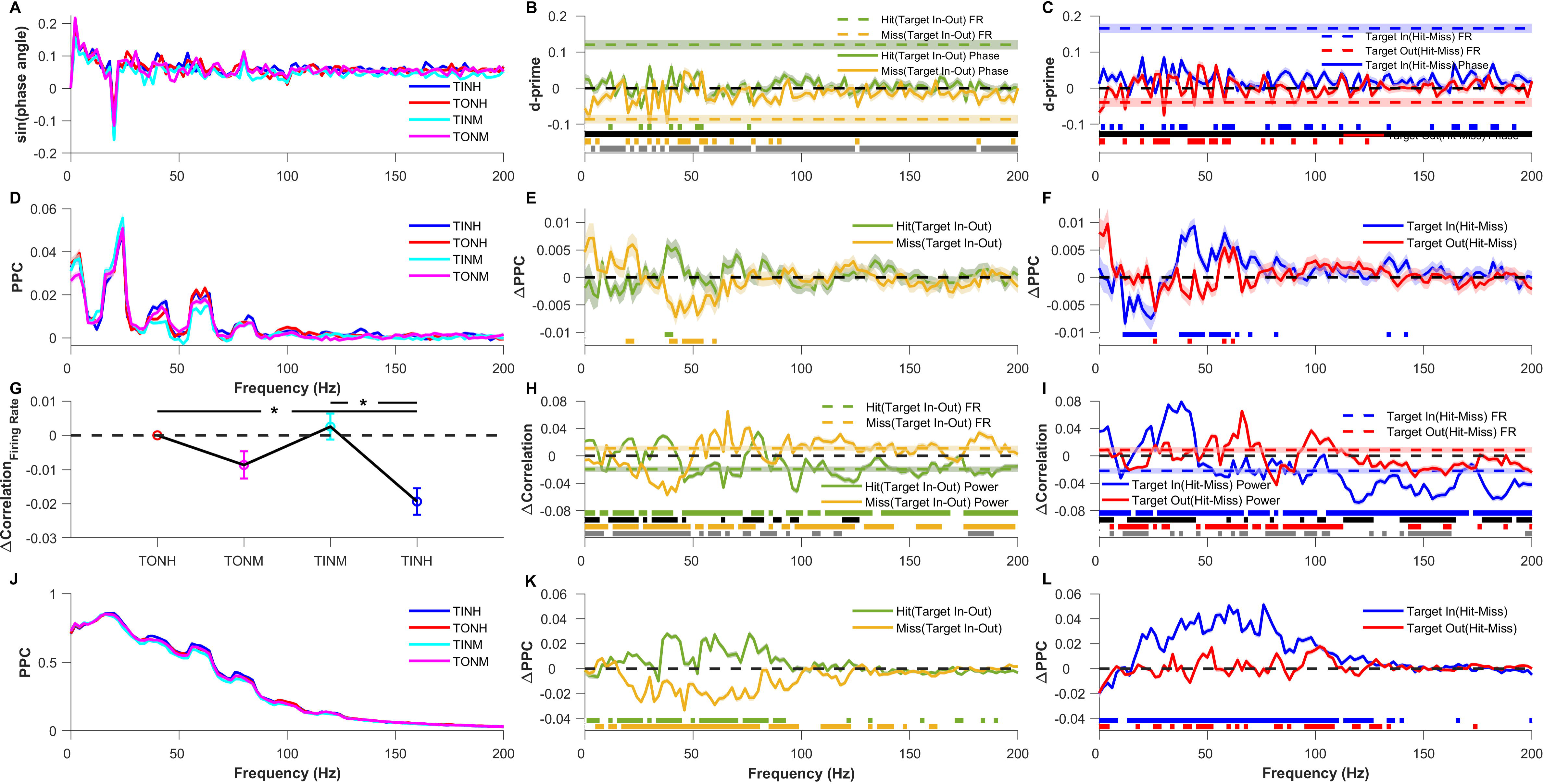
Comparison of LFP phase and pairwise phase consistency of individual electrodes and trial-wise firing rate correlation, LFP power correlation and pairwise phase consistency (PPC) of electrode pairs across neutrally cued attention and behavioral conditions. Same as figure 3 but for the neutrally cued conditions. In (A)–(F) mean and s.e.m are computed like in Supplementary Figure 1I – 1O. In (G)-(L) mean was taken across 50 bootstrap samples of mean across 5985 pairs and shaded lines and error bar indicate the bootstrap mean of s.e.m across 5985 electrode pairs. Horizontal color patches at the bottom of each panel indicate the significance level like in Figure 3.

**Supplementary Figure 3:**
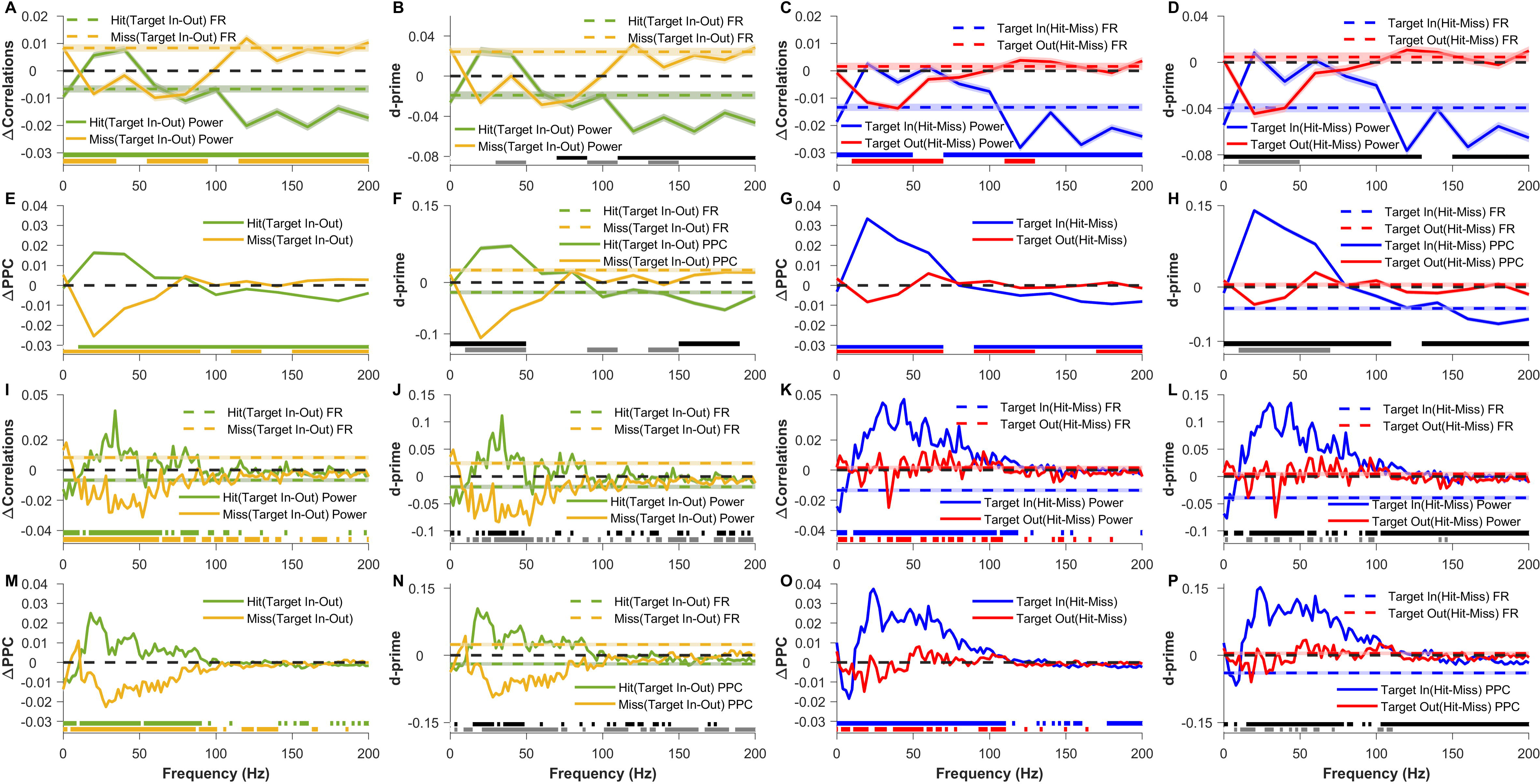
Comparison of single trial bin-wise firing rate correlation, bin-wise and taper-wise LFP power correlation and pairwise phase consistency (PPC) across neutrally cued attention and behavioral conditions. Related to Figure 5. Same as figure 5 but for neutrally cued conditions. Mean and s.e.m were computed like in Supplementary Figure 2G – 2L. Horizontal color patches at the bottom of each panel indicate the significance level like in Figure 5.

**Supplementary Figure 4:**
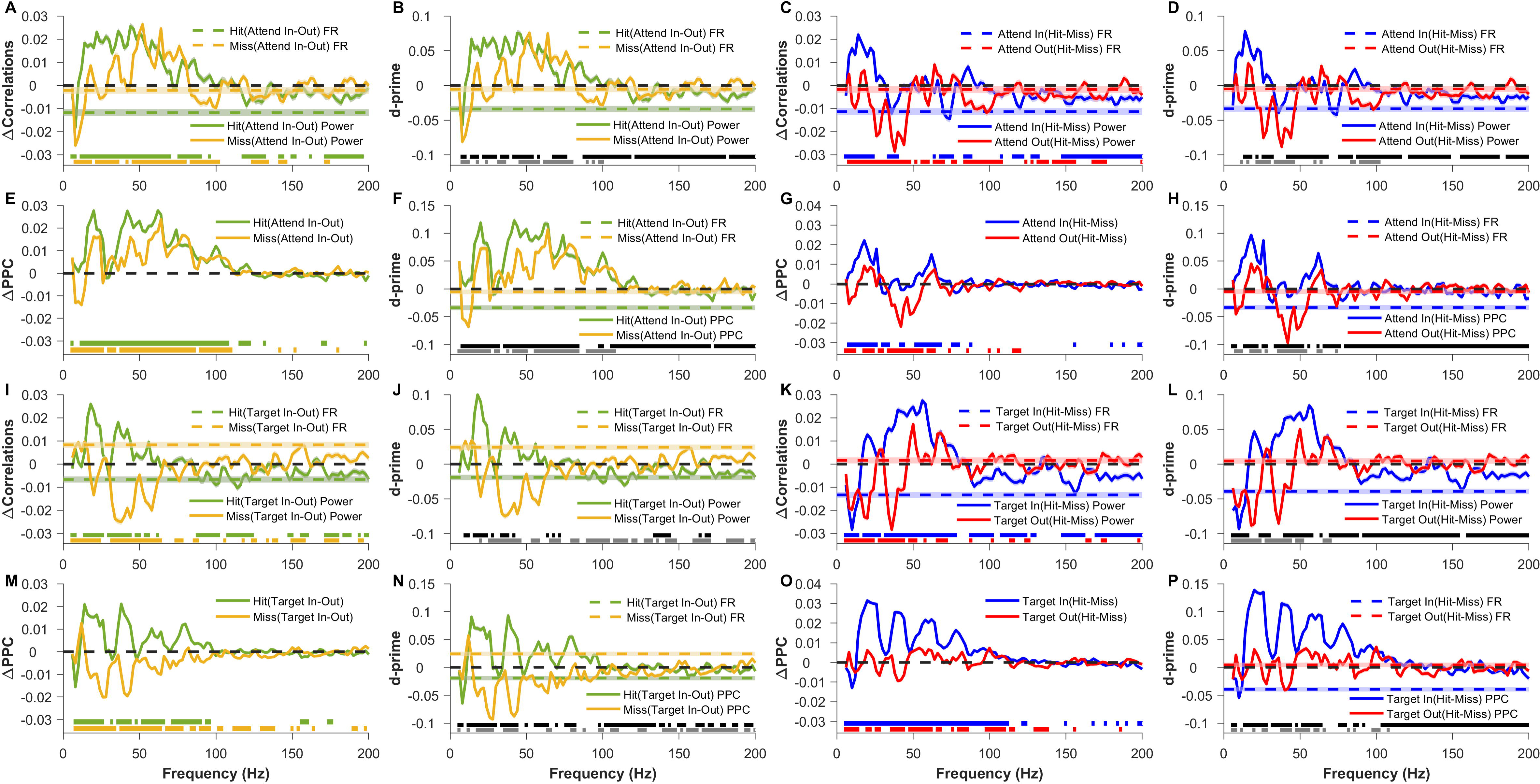
Comparison of single trial estimates of LFP power correlation and pairwise phase consistency (PPC) computed using Hilbert transform method with bin-wise firing rate correlation across valid and neutrally cued attention and behavioral conditions. (A)-(H) Same as figure 5I - P. Mean was taken across 50 bootstrap samples of mean across 5754 pairs and shaded lines indicate the bootstrap mean of s.e.m across 5754 electrode pairs. Horizontal color patches at the bottom of each panel indicate the significance level like in Figure 5. (I)-(P) Same as supplementary figure 3I-P. Mean was taken across 50 bootstrap samples of mean across 5985 pairs and shaded lines indicate the bootstrap mean of s.e.m across 5985 electrode pairs. Horizontal color patches at the bottom of each panel indicate the significance level like in (A)-(H).

**Supplementary Figure 5:**
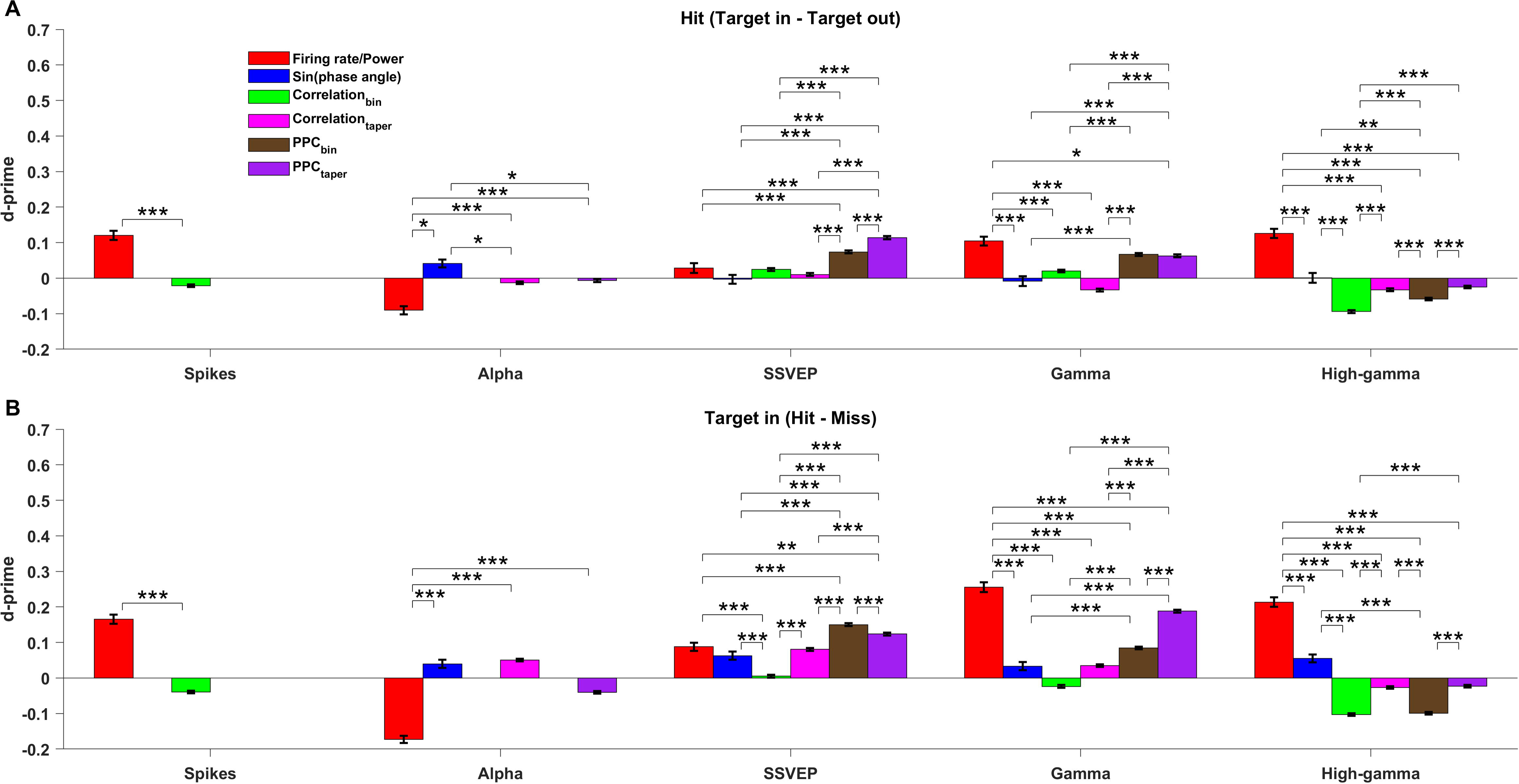
Summary plot comparing across the d-primes of all the measures for spikes, LFP power and phase in different frequency bands for neutrally cued attention and behavioral comparison. Same as figure 6 but for neutrally cued condition. Asterisks indicate the significance level like in Figure 6.

